# HMEC-1 extracellular vesicles as regulators of endothelial cell activation under inflammation

**DOI:** 10.64898/2026.06.08.730746

**Authors:** Ramon Castellanos-Sanchez, Scott Skalak, Matthew J. Lazzara, Luca Musante, Uta Erdbrügger, Shayn M. Peirce

## Abstract

Microvascular endothelial cell-derived extracellular vesicles (EVs) mediate local intercellular communication relevant to wound healing and inflammation, yet their proteomic cargo and functional properties remain poorly characterized. Here, EVs were isolated from human microvascular endothelial cells (HMEC-1) by standard ultracentrifugation (UC Bulk) or ultracentrifugation combined with size exclusion chromatography (UC+SEC) and characterized by nanoparticle tracking analysis, immunoblotting, cryogenic electron microscopy, and label-free mass spectrometry. UC+SEC achieved a 77-fold improvement in particle-to-protein ratio with 70–93% depletion of serum and extracellular matrix contaminants while preserving canonical EV markers (ALIX, CD9). Mass spectrometry identified 673 proteins in UC+SEC versus 336 in UC Bulk, with both preparations enriched in wound healing, hemostasis, and angiogenesis pathways. Despite dramatic purity differences, both isolation methods produced functionally comparable EVs that significantly enhanced dermal fibroblast wound closure. Functional assays on primary human dermal microvascular endothelial cells (HDMECs) revealed that HMEC-1-derived EVs exert inflammation-dependent dual effects on TNF-α pre-treated endothelium: upregulating VCAM-1 expression while simultaneously preserving VE-cadherin-mediated junction integrity. These effects were strictly inflammatory-dependent, with no detectable activity on healthy endothelial cells. This research uncovers a paradoxical phenotype in which microvascular endothelial EVs enhance immune cell recruitment signals while protecting barrier function exclusively under inflammatory conditions, suggesting a regulatory mechanism that may contribute to vascular homeostasis during inflammatory challenges.

## 1. INTRODUCTION

Extracellular vesicles (EVs) are membrane-enclosed nanoparticles (30–1000 nm) that mediate intercellular communication by transporting bioactive cargo including proteins, lipids, and nucleic acids between cells [1, 2]. These vesicles originate through diverse biogenesis pathways—either budding from the plasma membrane or forming within endosomal compartments before release—resulting in heterogeneous populations with distinct molecular compositions and biological functions [2, 3]. Regardless of their cellular origin, EVs coordinate biological processes across tissues by delivering their cargo to recipient cells through various mechanisms including membrane fusion, receptor-ligand interactions, and endocytic pathways [3].

The microvasculature—comprising arterioles, capillaries, and venules—constitutes approximately 80–90% of the vascular bed and represents the primary site of tissue-blood exchange [4, 5]. Microvascular endothelial cells form the innermost lining of these vessels and exhibit tissue-specific phenotypes distinct from their large vessel counterparts [6]. Beyond serving as a passive barrier, these specialized cells actively regulate critical vascular functions including tone, permeability, coagulation, inflammation, and angiogenesis [5, 7]. The unique microenvironment surrounding the microvasculature—containing pericytes, smooth muscle cells, immune cells, and tissue-specific parenchymal cells—creates complex signaling networks where local cell-to-cell communication plays essential roles in maintaining vascular homeostasis [8, 9]. Given these specialized functions and microenvironmental interactions, EVs released by microvascular endothelial cells likely contain unique cargo reflecting their distinct biological roles.

While EVs from large vessel endothelial cells (particularly HUVECs) and endothelial progenitor cells have been characterized in various contexts [10–12], significantly less attention has been directed toward microvascular endothelial cell-derived EVs. This knowledge gap is particularly relevant given the microvasculature’s central role in wound healing and inflammatory responses [13, 14]—processes requiring coordinated regulation of leukocyte recruitment, barrier function, and tissue regeneration through angiogenesis and paracrine signaling [15–17]. Understanding the composition and functional properties of microvascular endothelial cell-derived EVs may therefore provide insights into local vascular regulation during physiological and pathological conditions.

Current evidence from *in vitro* studies demonstrates that endothelial-derived EVs exert paracrine effects on neighboring vascular cells. Aortic endothelial cell-derived EVs promote inflammatory signaling in vascular smooth muscle cells through upregulation of adhesion molecules like VCAM-1 and transfer of pro-inflammatory proteins such as HMGB-1 and HMGB-2 [18]. The cargo composition and functional effects of endothelial-derived EVs are highly responsive to environmental cues, with factors like shear stress significantly altering their protein content [19]. However, the specific functional properties of microvascular endothelial cell-derived EVs—particularly their potential autocrine effects on endothelial cells regarding inflammatory regulation and barrier function—remain largely unexplored.

This study addresses these knowledge gaps through a multi-faceted approach. First, we compared two EV isolation methods—standard ultracentrifugation versus ultracentrifugation combined with size exclusion chromatography—to evaluate their impact on EV purity and functional activity. Second, we performed basic biophysical and biochemical characterization of the isolated EVs using nanoparticle tracking analysis, electron microscopy, and immunoblotting. Third, we conducted mass spectrometry analysis to characterize the proteomic cargo of microvascular endothelial cell-derived EVs. Finally, we evaluated the functional effects of these EVs in three cellular assays relevant to vascular homeostasis: wound healing in dermal fibroblasts, inflammatory marker expression in microvascular endothelial cells, and endothelial barrier integrity during inflammation. Our findings document the proteomic composition of these EVs and reveal context-dependent effects on inflammatory signaling and junction stability that may represent mechanisms for maintaining vascular function during inflammatory challenges.

## 2. METHODS

### 2.1 Cell culture

Human Microvascular Endothelial Cell Line-1 (HMEC-1, ATCC, Cat #CRL-3243) was cultivated in MCDB131 (Gibco, Cat #10372019) supplemented with recombinant human epidermal growth factor (rhEGF, 10 ng/mL), basic fibroblast growth factor (bFGF, 10 ng/mL), angiopoietin-1 (ANG1, 5 ng/mL), angiopoietin-2 (ANG2, 1 ng/mL), GlutaMAX (10 mM), hydrocortisone (1 µg/mL), ascorbic acid 2-phosphate (0.1 mM), HEPES buffer (5 mM), dibutyryl-cAMP (40.7 µM), IBMX (0.036 mM), heparin (25 µg/mL), AlbuMAX (0.5 µg/mL), and insulin-transferrin-selenium (ITS) mixture (500, 225, 0.335 mg/L, respectively); detailed reagent specifications including vendors, reconstitution protocols, and storage conditions are provided in Supplementary Table 1.

HMEC-1 cells were sequentially adapted to low-serum conditions by reducing fetal bovine serum (FBS, Gibco, Cat #A5256701) concentrations over three passages (10% to 5%, 5% to 2.5%, and 2.5% to 2%). All HMEC-1-derived EV collections were performed using cells between passages 6 and 12 cultured at 2% FBS.

For EV collection, HMEC-1 cells were seeded in T500 flasks at 500 cells/cm² in 100 mL of MCDB131 medium for 72 hours, washed once with 0.2 μm-filtered PBS, and maintained for 48 hours in MCDB131 medium supplemented with 2% EV-depleted FBS. EV depletion of FBS was performed by centrifugation at 2000 g for 20 minutes, filtration through 0.2 μm filter, and ultracentrifugation at 120,000 g for 144 minutes.

HMEC-1 cells were selected as the EV source due to their well-characterized microvascular endothelial phenotype, scalability for large-volume EV production, and reproducibility across passages.

All cells were cultured at 37°C with 5% CO□ in gelatin-coated flasks (0.1% gelatin in PBS).

### 2.2 EV isolation and separation

Eight independent EV isolation batches (ISO_1–ISO_8) were generated from HMEC-1 cells between passages 6 and 14 batch-specific details including cell seeding, NTA measurements, and experimental allocation are summarized in **Supplementary Table 2**.

#### 2.2.1 Ultracentrifugation (UC Bulk)

Conditioned medium was centrifuged at 2000 g for 20 minutes at 4°C to remove cells and large debris. The supernatant was filtered through a 0.45 μm filtration unit (VWR, Cat #10040) and subjected to differential ultracentrifugation: first at 9,266 rpm for 30 minutes to remove larger vesicles and cellular debris, followed by 32,000 rpm for 144 minutes to pellet small EVs, both steps performed using a Beckman Coulter 45Ti rotor (k-factor=262) in 70 mL polycarbonate tubes (Cat #355622). The resulting EV pellet was resuspended in 700 μL of 0.2 μm-filtered PBS. All centrifugation steps were performed at 4°C.

#### 2.2.2 Ultracentrifugation combined with size exclusion chromatography

For enhanced EV purification and removal of non-EV material including soluble proteins and protein aggregates, UC Bulk samples were further separated using size exclusion chromatography. An IZON qEV 70 nm second-generation column (Cat #ICO-70) was mounted on an IZON automatic fraction collector and prepared by saturation with 5 mL of 0.1% BSA in PBS, followed by extensive washing with 10 mL of 0.2 μm-filtered PBS (twice) and a final 15 mL PBS wash. The resuspended EV pellet (500 μL of the 700 μL UC Bulk total) was then loaded onto the prepared column. After a void volume of 2.90 mL, six sequential fractions of 1 mL each were collected.

Fraction 1 (FR1) was kept separate for characterization and functional assays and will be referenced as “UC+SEC” for simplicity though this article.

### 2.3 EV normalization and storage

For EV characterization by immunoblotting and mass spectrometry, protein normalization was performed using a Micro BCA™ Protein Assay Kit. For functional assays, nanoparticle tracking analysis (NTA) was used for normalization. UC Bulk samples (200 μL) were diluted to match NTA values of FR1, typically requiring 1:2 or 1:4 dilutions. All functional assays were conducted using EVs diluted 1:10 in MCDB131 medium.

For characterization and proteomic analysis, EV preparations were stored at 4°C for short-term use (24–120 hours) or at −80°C for long-term storage. For functional assays, EVs were stored exclusively at 4°C and used within one week of isolation.

### 2.4 EV characterization

#### 2.4.1 Nanoparticle tracking analysis

Particle size distribution and concentration were determined using a ZetaView system (Particle Metrix) with software version 8.05.12 SP. Samples were serially diluted to achieve concentrations between 5×10□ and 5×10□ particles/mL for optimal tracking. Video capture parameters included eleven measurement positions per sample with five measurement cycles, sensitivity 75, frame rate 30 frames/second, and shutter speed 75. All measurements were performed at room temperature after system calibration with 100 nm polystyrene beads.

#### 2.4.2 Immunoblotting

Both EV preparations were characterized for canonical EV markers ALIX (Thermo Cat#3A9) and CD9 (BioLegend Cat# 312102). Given their high protein content UC Bulk samples were directly loaded into gels while UC+SEC fraction samples were concentrated using trichloroacetic acid precipitation (0.4%) with 4% sodium deoxycholate and resuspended in 2× Laemmli buffer. Equal protein amounts (10 μg) were loaded per lane on 4–12% Bis-Tris polyacrylamide gels. After electrophoresis, proteins were transferred to nitrocellulose membranes. Membranes were blocked for 1 hour in Intercept Blocking Buffer (LI-COR Biosciences) and incubated with primary antibodies overnight at 4°C. Primary antibodies were used at 1:1000 dilution. Secondary antibodies included IRDye 800CW and IRDye 680RD (LI-COR Biosciences, 1:5,000). Imaging was performed on an Odyssey Infrared Imager (LI-COR Biosciences). Full nitrocellulose membrane detection is presented in Supplementary Figure 1.

#### 2.4.3 Cryo-electron microscopy

EV morphology was examined using cryogenic electron microscopy at University of Virginia Electron Microscopy Core. Samples were prepared on Quantifoil R2/2 holey carbon copper grids (200 mesh) after glow-discharge treatment (45 seconds at 15 mA). A Vitrobot Mark IV (FEI/Thermo Fisher) was used for sample vitrification with the following parameters: 4°C chamber temperature, 100% humidity, blot force −2, blot time 4 seconds. Grids were rapidly plunged into liquid ethane, transferred to grid boxes under liquid nitrogen, and stored until imaging. Data were acquired using a 300kV FEI Titan Krios transmission electron microscope at 75,000× magnification (1.08 Å/pixel) with a total electron dose limited to 32 e□/Å² to minimize radiation damage. Images were collected across a defocus range from −1.0 to −3.0 μm to enhance phase contrast for optimal visualization of EV membranes.

### 2.5 Mass spectrometry analysis

#### 2.5.1 Sample preparation

EV samples containing 7–8 μg total protein were resuspended in 5% SDS buffer containing 50 mM triethylammonium bicarbonate (TEABC) and 20 mM DTT, and heated for 10 minutes at 95°C. After cooling to room temperature, iodoacetamide in 5% SDS solution was added to a final concentration of 40 mM and incubated in the dark for 30 minutes. Samples were centrifuged for 8 minutes at 13,000 g, and supernatants were processed using S-Trap columns (ProtiFi, LLC). Proteins were digested with sequencing-grade Lys-C/trypsin (Promega) by overnight incubation at 37°C. The resulting peptides were eluted and dried using a SpeedVac (Fisher Scientific).

#### 2.5.2 NanoUPLC-MS/MS analysis

Peptides were analyzed using a nanoAcquity UPLC system (Waters) coupled with an Orbitrap Fusion Lumos mass spectrometer (Thermo Fisher) at Georgetown’s University Medical Center Mass Spectrometry and Analytical Pharmacology Core. Samples in 0.1% formic acid solution were loaded onto a C18 trap column (Waters Acquity UPLC M-Class Trap, Symmetry C18, 100 Å, 5 μm, 180 μm × 20 mm) at 10 μL/min for 4 minutes. Peptides were then separated on an analytical column (Waters Acquity UPLC M-Class, peptide BEH C18 column, 300 Å, 1.7 μm, 75 μm × 150 mm) maintained at 45°C with a flow rate of 350 nL/min. Separation was achieved using a 150-minute gradient of buffer A (2% ACN, 0.1% formic acid) and buffer B (0.1% formic acid in ACN): 1% buffer B at 0 min, 5% buffer B at 1 min, 22% buffer B at 90 min, 50% buffer B at 100 min, 98% buffer B at 120 min, 98% buffer B at 130 min, 1% buffer B at 130.1 min, and 1% buffer B at 150 min.

The mass spectrometer was operated with an ion spray voltage of 2.2 kV and an ion transfer temperature of 275°C. MS parameters included Orbitrap resolution 60,000, scan range 380–1400 m/z, RF lens 30%, standard AGC target, and automatic maximum injection time. MS/MS parameters included quadrupole isolation with 1.6 m/z isolation window, HCD activation at 35% collision energy, Orbitrap detection at 30,000 resolution, and normalized AGC target of 200%.

#### 2.5.3 Data analysis

MS data files were processed using Proteome Discoverer platform (version 2.4, Thermo Scientific) with the Sequest HT algorithm. MS/MS spectra were searched against the human proteome database with parameters including two allowed missed cleavages, minimum peptide length of seven amino acids, oxidation (M) as variable modification, carbamidomethylation (C) as fixed modification, and MS and MS/MS ion tolerances of 10 ppm and 0.02 Da, respectively. False discovery rate (FDR) was estimated using fixed value PSM validation. For comparative analysis between samples, total unique peptide counts with a minimum threshold of three peptides were used to identify proteins. For each sample, protein intensities were normalized to GAPDH; normalized values were then converted to z-scores within each sample (column-wise) across the displayed proteins. Gene Ontology (GO) term analysis was performed using the clusterProfiler package in R (version 2023.03.0+386).

### 2.6 Endothelial activation assay

To analyze EV effects on endothelial activation, two distinct models were employed using human Dermal Microvascular Endothelial Cells (HDMEC, PromoCell, Cat# C-12210) cultured in MCDB131 medium supplemented with 2% FBS and seeded at 20,000 cells/cm². HDMECs were used as recipient cells in functional assays to ensure physiological relevance, as primary cells more closely recapitulate *in vivo* endothelial responses to inflammatory stimuli and EV-mediated signaling than immortalized cell lines.

For the inflamed model, cells were adapted to serum-free conditions for 6 hours before TNF-α treatment (10 ng/mL) for 16 hours to induce inflammatory activation. For the healthy model, PBS was substituted for TNF-α during pre-treatment. Following pre-treatment, medium was replaced with fresh medium containing EV preparations or PBS control, and cells were incubated for 24 hours before analysis.

The experimental groups included TNF-α or PBS control (pre-treated, no EV treatment), UC Bulk, and UC+SEC treatments. EV stock preparations (5–8×10¹□ particles/mL) were diluted 1:10 in MCDB131 medium, with 1.2 mL added per well, corresponding to approximately 6–9.6×10□ nanoparticles per well or 60–96×10³ nanoparticles per cell. Experiments were performed in technical triplicates using a single batch of primary HDMECs, with inflamed and healthy models conducted using separate EV batches.

For immunocytochemistry analysis, cells were fixed with 4% paraformaldehyde for 20 minutes, permeabilized with 0.1% Triton X-100 for 5 minutes and blocked with Odyssey blocking buffer for 1 hour. Primary antibodies against VE-cadherin (Thermo Cat#16B1, 1:400) and VCAM-1 (Abcam Cat#EPR5047, 1:200) were applied overnight at 4°C, followed by secondary antibodies (Alexa Fluor 488 and 594, 1:500) for 1 hour and DAPI counterstaining. Intercellular gap quantification was performed by measuring VE-cadherin-negative areas between adjacent cells using ImageJ software. For immunoblotting analysis, cells were lysed in RIPA buffer supplemented with protease inhibitors, and protein expression was assessed using the same primary antibodies with IRDye secondary antibodies and Odyssey infrared imaging. Full nitrocellulose membrane detection is presented in Supplementary Figure 2.

### 2.7 Wound healing assay

EVs effects on wound healing were tested on Human Dermal Fibroblasts adult (HDFa, ATCC Cat# PCS-201-012). HDFa were seeded at 100,000 cells per well in 48-well plates in Fibroblast Basal Medium (ATCC Cat# PCS-201-030) supplemented with Low-serum growth kit (ATCC Cat# PCS-201-041) and cultured for 24 hours until confluence.

A standardized scratch wound was created in confluent monolayers using a 200 μL pipette tip, with precise locations marked using the position-saving feature of a Leica Thunder microscope. After washing with PBS to remove debris, culture medium was replaced with EV-containing medium prepared by diluting basal medium with EV preparations 1:10. Experimental groups included PBS control, UC Bulk, and UC+SEC treatments. EVs were applied at 7×10□ nanoparticles/mL (2.1×10□ total nanoparticles per well, 21×10³ nanoparticles per cell).

Following 16-hour incubation, cells were fixed with 4% paraformaldehyde for 20 minutes, permeabilized with 0.1% Triton-X for 5 minutes, blocked with Odyssey blocking buffer for 1 hour, and stained with phalloidin (Invitrogen Cat# A30107) for 1 hour to visualize F-actin cytoskeleton. Images were captured at 0 and 16 hours at identical positions using stored coordinates. Wound closure was quantified using ImageJ software by converting phalloidin-stained images to 8-bit format and calculating the percentage of wound closure as: (1 – (Final wound area / Initial wound area)) × 100. All experiments were performed in biological triplicates with three technical replicates per condition.

### 2.8 Statistical analysis

Statistical analyses were performed using GraphPad Prism 10 (Version 10.2.3, GraphPad Software Inc.). For multiple group comparisons, one-way analysis of variance (ANOVA) was employed, followed by Tukey’s honestly significant difference (HSD) post-hoc test. Each experimental model (inflamed/healthy endothelium, fibroblasts) was analyzed independently using data from three technical replicates per condition. For correlation analyses of proteomic data, Pearson’s correlation coefficient (r) was calculated to assess relationships between protein expression profiles across different EV isolation methods. Proteins with less than three unique peptides were excluded from correlation analysis to ensure reliability. Differences with p-values < 0.05 were considered statistically significant. All results are expressed as mean ± standard deviation.

## 3. RESULTS

### 3.1 Comparative characterization of EV isolation methods

We compared two EV isolation approaches: standard ultracentrifugation (UC Bulk, representing the EV pellet obtained by differential centrifugation with 0.45 μm pre-filtration, resuspended in PBS) and ultracentrifugation followed by size exclusion chromatography (UC+SEC) (Figure 1A).

**Figure 1.**
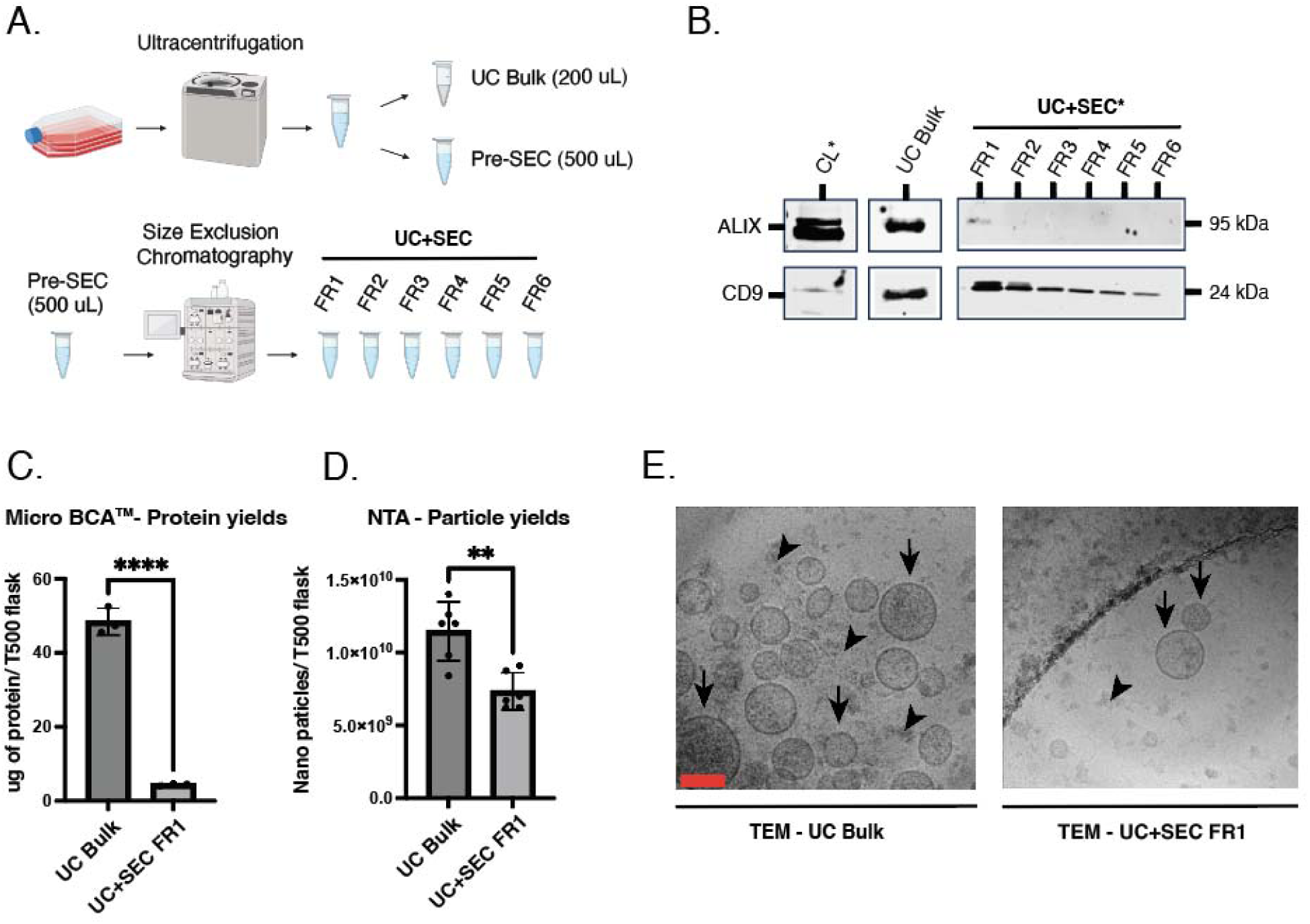
Comparative characterization of EV isolation methods. (A) Schematic overview of EV isolation workflow comparing ultracentrifugation (UC Bulk) and ultracentrifugation combined with size exclusion chromatography for Fraction 1 (UC+SEC FR1). (B) Immunoblot analysis of canonical EV markers ALIX and CD9. (*) The cell Lysate (CL) and UC+SEC fractions were TCA/DOC reduced. (C) Protein yield quantification by Micro-BCA assay. (D) Nanoparticle tracking analysis of particle yield. (E) Cryogenic electron microscopy of UC Bulk and UC+SEC FR1 preparations, scale 50 nm. Arrows indicate intact vesicles with defined lipid bilayers; arrowheads indicate non-vesicular background material. Statistical significance at ****p < 0.0001, **p < 0.01. Full uncropped membranes are provided in Supplementary Figure 1.

#### 3.1.1 EV marker distribution and protein content

Immunoblot analysis showed similar enrichment of canonical EV markers between isolation methods Both UC Bulk and UC+SEC preparations demonstrated positivity for established EV markers ALIX and CD9. For microvascular endothelial cells derived-EVs, the western blots results indicate that CD9 was more enriched in EV preparations than ALIX in comparison with the cell lysate (CL). (Figure 1B).

Protein quantification showed differences between methods (Figure 1C). UC Bulk preparations contained an average of 48 μg protein per T500 flask (100 mL culture media processed), while UC+SEC yielded 4.4 μg protein per T500 flask, representing a 91% reduction in total protein content. Protein quantification was performed using the Micro-BCA assay with samples normalized by total protein content (μg) for immunoblot analysis and mass spectrometry.

#### 3.1.2 Particle yield

Nanoparticle tracking analysis showed differences in yield (Figure 1D). UC Bulk preparations yielded 1.03±0.8×10¹□ particles per T500 flask, while UC+SEC contained 7.3±0.2×10□ particles per T500 flask, representing a 29% reduction in particle count.

Particle-to-protein ratio calculations showed differences between isolation methods. UC Bulk showed 2.15×10□ particles/μg protein, while UC+SEC demonstrated a markedly higher ratio of 1.66×10¹□ particles/μg protein, representing a 77-fold increase, indicating a significant lower degree of non-EV proteins in the UC+SEC preparations.

#### 3.1.3 Ultrastructural characterization by cryogenic electron microscopy

Cryogenic electron microscopy confirmed the presence of intact vesicular structures in both UC Bulk and UC+SEC preparations (Figure 1E). UC+SEC samples exhibited substantially reduced non-vesicular background debris compared to UC Bulk preparations (Figure 1E arrowheads). The UC+SEC preparation contained predominantly intact vesicles with clearly defined lipid bilayers ranging from 30–150 nm in diameter. UC Bulk preparations showed greater heterogeneity, including vesicle aggregates and non-vesicular material.

### 3.2 Mass spectrometry analysis

Mass spectrometry analysis was performed to characterize proteomic profiles of UC Bulk and UC+SEC preparations, with a minimum detection threshold of three unique peptides across technical triplicates. Venn diagram analysis showed that UC+SEC detected 673 proteins compared to 336 proteins in UC Bulk, with 299 proteins shared between both methods. Comparison with the Vesiclepedia Top 100 database of most frequently reported EV proteins showed that UC+SEC identified 96 of these proteins (83 detected by both methods plus 13 detected exclusively by UC+SEC), while UC Bulk detected 83 proteins (Figure 2A). Complete mass spectrometry data including unique peptide counts for all identified proteins across UC Bulk and UC+SEC triplicates are provided in Supplementary Table 3.

**Figure 2.**
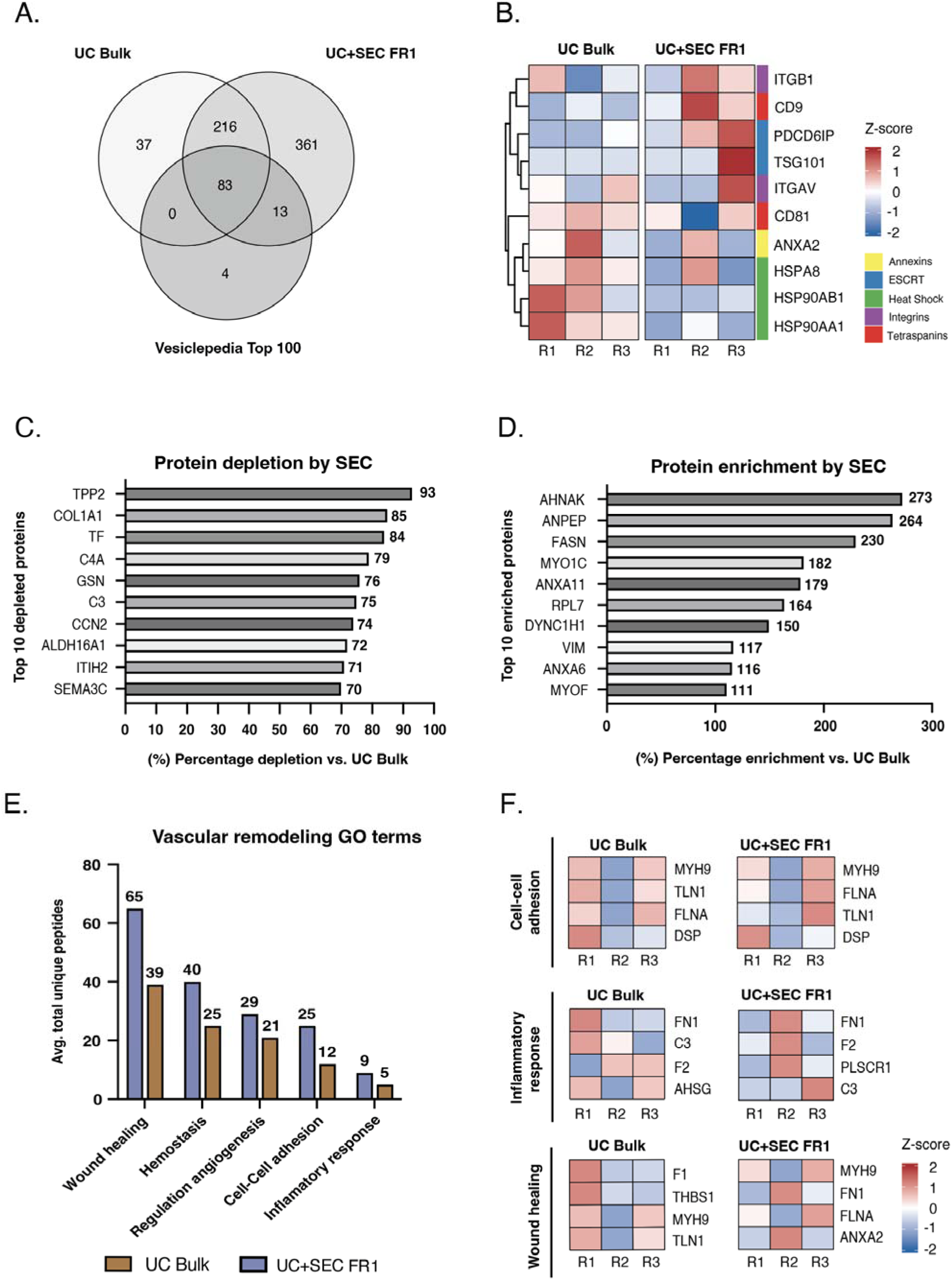
Mass spectrometry proteomic analysis of UC Bulk and UC+SEC FR1 preparations. (A) Venn diagram of protein identification overlap. (B) Heatmap of GAPDH-normalized expression of canonical EV markers. (C) Top depleted proteins in UC+SEC FR1 versus UC Bulk. (D) Top enriched proteins in UC+SEC FR1 versus UC Bulk. (E) Gene Ontology biological process enrichment analysis. (F) Top abundant proteins per GO term category.

Figure 2B displays GAPDH-normalized expression profiles of 10 representative canonical EV markers, including the ESCRT components TSG101 and ALIX/PDCD6IP. Values are GAPDH-normalized and z-scored within each sample (column-wise), so the heatmap reflects the relative ranking of markers within a given preparation rather than a direct comparison of individual markers between methods. Mass spectrometry analysis detected 16 of 18 MISEV-recommended markers (88.9%) across both isolation methods. Of these 16 markers, 12 were detected in both UC Bulk and UC+SEC preparations, while 4 markers (TSG101, FLOT1, FLOT2, RAB5A) were exclusively identified in UC+SEC samples.

Among the 12 markers detected by both methods, expression levels showed consistency, with 10 of 12 markers (83.3%) displaying similar abundance (0.7–1.4× fold change range). Average GAPDH-normalized expression was nearly identical between methods (UC Bulk = 0.770, UC+SEC = 0.757; 0.98× overall fold change). The most abundant markers displayed in the heatmap—ANXA2, HSPA8, and heat shock proteins HSP90AA1/HSP90AB1—showed robust detection across all replicates in both methods. Average z-scores across all 16 detected markers were balanced between methods (UC Bulk = 0.09, UC+SEC = −0.09).

Analysis of UC+SEC versus UC Bulk proteomes showed depletion of proteins associated with non-EV contamination (Figure 2C). Twenty-five proteins showed greater than 50% depletion in UC+SEC compared to UC Bulk, with the top 10 most depleted proteins displaying 70–93% reduction. These highly depleted proteins comprised three major categories: serum-derived proteins including transferrin (TF, 84% depletion), complement components C3 and C4A (75% and 79%, respectively), plasma gelsolin (GSN, 76%), and inter-alpha-trypsin inhibitor heavy chain 2 (ITIH2, 71%); extracellular matrix proteins including collagen type I alpha 1 (COL1A1, 85%), connective tissue growth factor (CCN2, 74%), and semaphorin 3C (SEMA3C, 70%); and intracellular proteins including tripeptidyl peptidase II (TPP2, 93%) and aldehyde dehydrogenase 16A1 (ALDH16A1, 72%). None of the depleted proteins are listed as EV markers in MISEV guidelines.

Analysis showed 163 proteins with >20% enrichment in UC+SEC compared to UC Bulk (Figure 2D). Proteins were selected using dual criteria of fold change >1.5× and robust detection (≥10 unique peptides). The top 10 proteins meeting these criteria exhibited 111–273% enrichment in UC+SEC and comprised several functional categories, including membrane remodeling proteins (ANXA11, ANXA6, MYOF), cytoskeletal scaffolding components (AHNAK, VIM, DYNC1H1, MYO1C), the endothelial marker ANPEP (CD13), and fatty acid synthase (FASN).

Gene Ontology enrichment analysis showed that both UC Bulk and UC+SEC preparations were enriched in biological processes related to vascular remodeling and tissue repair (Figure 2E). Wound healing was the most enriched category, with 65 proteins in UC+SEC and 39 proteins in UC Bulk. Hemostasis showed enrichment with 40 proteins in UC+SEC and 25 proteins in UC Bulk, while regulation of angiogenesis showed 29 proteins in UC+SEC and 21 proteins in UC Bulk. Cell-cell adhesion and inflammatory response categories contained 25 and 9 proteins in UC+SEC, compared to 12 and 5 proteins in UC Bulk, respectively. UC+SEC detected more proteins across all functional categories compared to UC Bulk, while both preparations showed enrichment in the same vascular remodeling pathways.

The top four most abundant proteins within each GO term category were analyzed for both preparations (Figure 2F). For cell-cell adhesion, both methods showed enrichment of cytoskeletal proteins including MYH9 (non-muscle myosin IIA), FLNA (filamin A), TLN1 (talin-1), and DSP (desmoplakin). The inflammatory response category showed enrichment of fibronectin (FN1), complement component C3, and coagulation factor II (F2) in both methods, with UC Bulk additionally containing AHSG (fetuin-A) and UC+SEC showing enrichment of PLSCR1 (phospholipid scramblase 1). Wound healing proteins showed greater compositional differences: UC Bulk showed high enrichment of THBS1 (thrombospondin-1), while UC+SEC showed greater enrichment of ANXA2 (annexin A2), though both preparations shared FN1 and MYH9 as abundant wound healing proteins. Despite these compositional differences in specific proteins, both UC Bulk and UC+SEC showed enrichment across the same functional categories.

### 3.3 Inflammation-dependent effects of EVs on endothelial activation and barrier function

Based on proteomic identification of cell adhesion proteins (Integrins and cadherins) and inflammatory response proteins (10 proteins, adjusted p-value < 0.057) in HMEC-1-derived EVs, we investigated their functional effects on endothelial activation using primary human dermal microvascular endothelial cells (HDMECs). Two experimental models were employed: a healthy model (PBS pre-treatment) and an inflamed model (TNF-α pre-treatment at 10 ng/mL for 16 hours). Following pre-treatment, cells were cultured with EV preparations (approximately 6–9.6×10□ nanoparticles per well; 60–96×10³ nanoparticles per cell) or PBS control for 24 hours. Immunofluorescence and immunoblot analyses revealed no significant changes in VCAM-1 expression in healthy conditions across all EV treatments (Figure 3A, 3C and 3D). In contrast, TNF-α pre-treated cells exhibited marked upregulation of VCAM-1 following EV treatment (Figure 3B, 3E and 3F). UC+SEC treatment demonstrated the most pronounced effect, with a 2.0-fold increase in VCAM-1/GAPDH ratio compared to TNF-α controls (p<0.05, Figure 3F). UC Bulk treatment showed a 1.6-fold increase in VCAM-1 expression. VE-cadherin expression remained stable across all conditions in both healthy and inflamed models (Figure 3D, 3F). Evaluation of endothelial barrier integrity revealed differential effects dependent on inflammatory context. In healthy conditions, VE-cadherin staining showed continuous junctional localization with minimal intercellular gaps across all conditions, averaging 2.0±0.5 μm² per field with no significant differences between groups (Figure 3G, 3H). In TNF-α pre-treated cells, substantial junctional disruption was observed in control conditions, with intercellular gap areas averaging 25.0±2.5 μm² per field (Figure 3G, 3H). EV treatment significantly reduced these gaps: UC Bulk treatment decreased gap areas to 3.5±1.0 μm² (p<0.01), while UC+SEC treatment reduced gaps to 6.0±1.5 μm² (p<0.05) compared to TNF-α controls (Figure 3H). These data demonstrate that HMEC-1-derived EVs exert inflammation-specific effects on endothelial activation, simultaneously enhancing VCAM-1 expression while preserving VE-cadherin-mediated junction integrity exclusively in TNF-α pre-treated endothelium.

**Figure 3.**
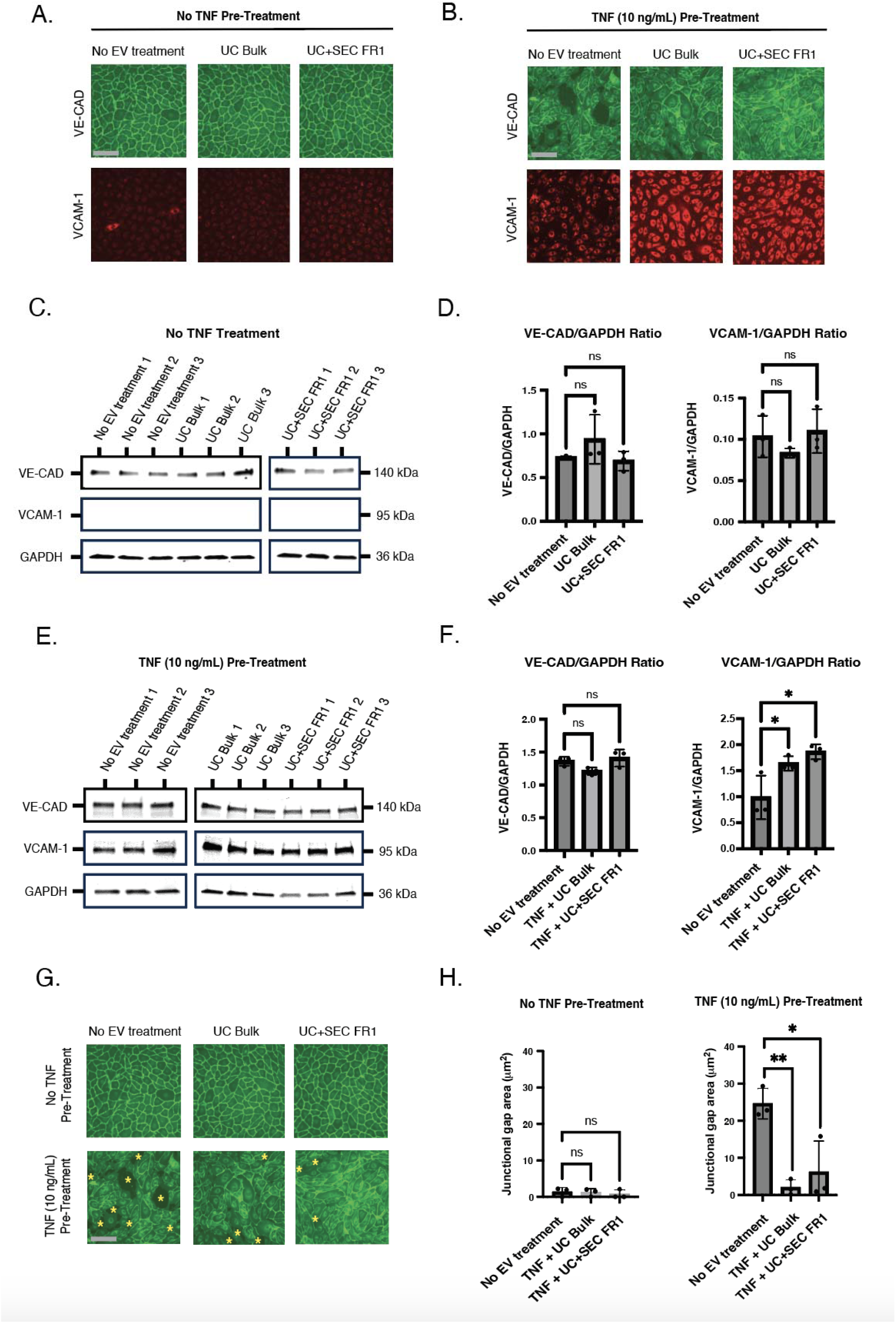
HMEC-1-derived EVs exert inflammation-dependent effects on endothelial activation and barrier function. (A) Representative immunofluorescence images of primary human dermal microvascular endothelial cells (HDMECs) showing VE-cadherin (VE-CAD, green) and VCAM-1 (red) expression in healthy conditions (no TNF-α pre-treatment) following treatment with PBS control, UC Bulk EVs, or UC+SEC FR1 EVs. (B) Representative immunofluorescence images of HDMECs showing VE-CAD (green) and VCAM-1 (red) expression in inflamed conditions (TNF-α 10 ng/mL pre-treatment for 16 hours) following EV treatments. (C) Western blot analysis of VE-CAD, VCAM-1, and GAPDH expression in HDMECs under healthy conditions across three biological replicates for each treatment. (D) Quantification of VE-CAD/GAPDH and VCAM-1/GAPDH ratios in healthy conditions (n = 3). (E) Western blot analysis of VE-CAD, VCAM-1, and GAPDH expression in TNF-α pre-treated HDMECs across three biological replicates for each treatment. (F) Quantification of VE-CAD/GAPDH and VCAM-1/GAPDH ratios in TNF-α pre-treated conditions (n = 3). (G) Representative immunofluorescence images showing VE-CAD junctional localization (green) in both healthy and TNF-α pre-treated conditions. Yellow asterisks indicate intercellular gaps in TNF-α treated controls. Scale bar = 50 μm. (H) Quantification of junctional gap areas in healthy and TNF-α pre-treated HDMECs following EV treatments (n = 3). ns = not significant, *p < 0.05, **p < 0.01. Full uncropped membranes are provided in Supplementary Figure 2.

### 3.4 Effects of HMEC-1-derived EVs on dermal fibroblast wound healing

To assess the wound healing potential of HMEC-1-derived EVs, scratch wound assays were performed on monolayer cultures of human dermal fibroblasts (HDFs). Cells were seeded at 100,000 cells/well in 48-well plates 24 hours prior to the assay. Following wound creation, cells were treated with normalized EV doses (2.1×10□ total particles per well; 21×10³ particles/cell) or PBS vehicle control, and migration was assessed after 16 hours.

Phase contrast microscopy showed wound closure in all conditions (Figure 4A, top panels). Phalloidin staining revealed reorganized actin cytoskeleton at the wound edge in EV-treated fibroblasts, with actin filaments in the direction of migration (Figure 4A, middle panels). These cytoskeletal changes were most pronounced in UC+SEC-treated cells.

**Figure 4.**
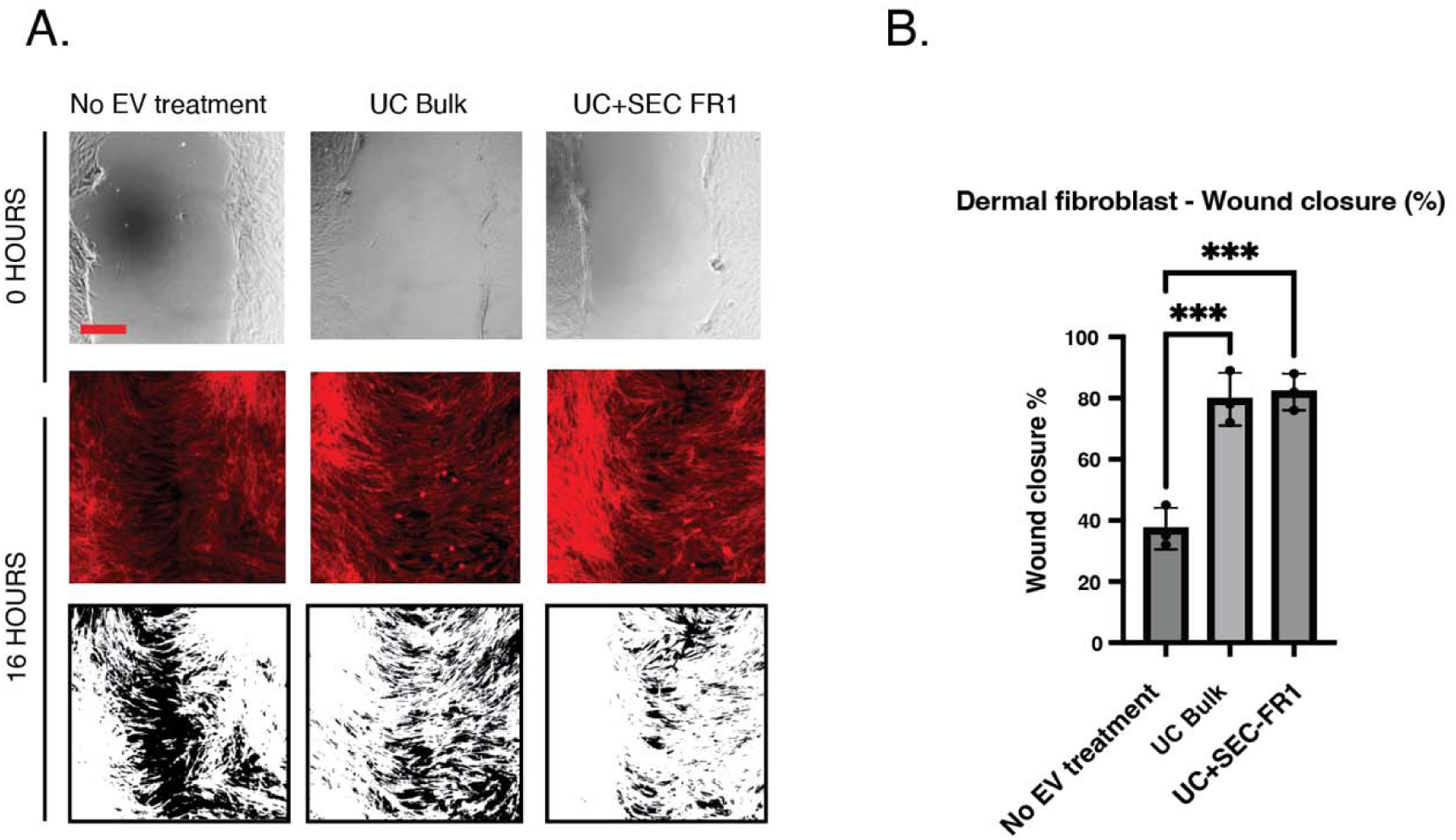
Effects of HMEC-1-derived EVs on dermal fibroblast wound healing. (A) Representative phase contrast images (top panels), phalloidin staining showing actin cytoskeleton organization (middle panels, red), and processed wound edge images (bottom panels, black and white) of human dermal fibroblasts at 0 hours and 16 hours following scratch wound creation and treatment with PBS control (No EV treatment), UC Bulk, or UC+SEC FR1. Scale bar = 200 μm. (B) Quantification of wound closure percentage at 16 hours (n = 3). ***p < 0.001.

Quantitative analysis showed that all EV preparations significantly enhanced wound closure compared to PBS control (Figure 4B). Control conditions achieved 37±4% wound closure, while UC Bulk treatment resulted in 79±6% closure and UC+SEC treatment achieved 82±5% closure (p<0.001 versus control for both EV treatments). These results demonstrate that HMEC-1-derived EVs significantly enhance dermal fibroblast migration and wound closure.

## 4. DISCUSSION

### 4.1 SEC effectively removes protein contaminants while preserving EV integrity

Our finding that UC+SEC achieved a particle-to-protein ratio of 1.66×10¹□ particles/μg protein compared to 2.15×10□ for UC Bulk confirms and extends previous reports demonstrating SEC’s superior particle-to-protein ratio [20,21,22]. The 77-fold improvement aligns with established quality control metrics, where ratios approaching 3×10¹□ particles/μg indicate minimal protein contamination [23,24].

The specific protein depletion profile we observed with 70–93% reduction in serum-derived proteins (transferrin, complement C3/C4A, gelsolin), extracellular matrix proteins (COL1A1, CCN2, SEMA3C), and intracellular contaminants (TPP2, ALDH16A1) validated SEC’s capacity to remove major non-EV protein classes that confound ultracentrifugation preparations [25,26]. The 85% depletion of COL1A1 and substantial reduction of fibronectin are particularly notable, as these ECM proteins represent persistent contaminants in ultracentrifugation-only protocols [27,28].

Critically, both UC Bulk and UC+SEC preparations showed similar enrichment of canonical EV markers by Western blot (ALIX, CD9), and mass spectrometry detected 16 of 18 MISEV-recommended markers (88.9%) with balanced z-scores between methods (UC Bulk = 0.09, UC+SEC = −0.09). This indicates that SEC purification retains the core EV protein complement while selectively removing co-isolated contaminants, a crucial distinction from purification methods that achieve high purity by sacrificing EV recovery.

### 4.2 Proteomic characterization reveals wound healing and hemostasis pathway enrichment

Mass spectrometry analysis demonstrated that UC+SEC detected 673 proteins compared to 336 in UC Bulk, with 299 proteins shared between methods. The enhanced protein identification in SEC-purified samples likely reflects removal of abundant contaminating proteins that suppress ionization of lower-abundance EV proteins during mass spectrometry [33,34]. This represents a technical advantage beyond purity metrics—SEC enables more comprehensive proteomic characterization of EV cargo.

Gene Ontology enrichment analysis revealed that both UC Bulk and UC+SEC preparations were enriched in biological processes related to vascular remodeling and tissue repair, with wound healing being the most enriched category (65 proteins in UC+SEC vs 39 proteins in UC Bulk). Hemostasis showed enrichment with 40 proteins in UC+SEC versus 25 proteins in UC Bulk, while regulation of angiogenesis contained 29 versus 21 proteins, respectively. These pathway enrichments align with extensive literature documenting MSC and cardiac progenitor cell EV proteomes, which consistently identify hemostasis, ECM organization, complement cascades, and platelet degranulation pathways [35].

The top four most abundant proteins within each GO term category showed interesting compositional differences. For wound healing, UC Bulk showed high enrichment of thrombospondin-1 (THBS1), while UC+SEC showed greater enrichment of annexin A2 (ANXA2), though both preparations shared fibronectin (FN1) and non-muscle myosin IIA (MYH9). These compositional differences may explain subtle functional variations that did not reach significance in our assays but could emerge in more sensitive or mechanistically targeted experiments.

Importantly, despite these compositional differences in specific proteins, both UC Bulk and UC+SEC showed enrichment across the same functional categories, suggesting both preparations retain the core wound healing and angiogenesis-promoting cargo. This functional pathway conservation despite protein-level differences provides mechanistic rationale for the comparable bioactivities we observed and indicates that multiple protein combinations can mediate similar biological outcomes—a concept of redundancy common in complex biological systems.

### 4.3 Both isolation methods preserve functional activity despite dramatic purity differences

A central and somewhat unexpected finding is that UC Bulk and UC+SEC preparations exhibited comparable functional activities in wound healing assays (79±6% vs 82±5% closure, respectively) and endothelial barrier protection (gap reduction to 3.5±1.0 μm² vs 6.0±1.5 μm²) despite the 77-fold difference in particle-to-protein ratios. This challenges the implicit assumption that higher purity necessarily correlates with enhanced bioactivity and suggests that co-isolated proteins in UC Bulk preparations do not substantially interfere with core EV-mediated functions in these specific assays. However, as Whittaker et al. [29] demonstrated, soluble factors co-isolated with EVs can contribute to observed bioactivity, and the possibility that co-purified proteins partially contribute to the functional effects in UC Bulk preparations cannot be excluded.

Recent work by Fernández-Sáez et al. [30] provides important context: while they similarly reported superior particle-to-protein ratios for SEC-EVs (1.13×10□) versus UC-EVs (7.83×10□), they observed functional divergence—SEC-EVs enhanced platelet aggregation by 16.67±4.20% (p=0.0005) while UC-EVs showed no effect. This contrasts with our findings where both preparations remained functionally active. The discrepancy likely reflects differences in functional readout sensitivity and the specific biological processes examined—platelet aggregation may require higher purity EVs than endothelial activation or wound healing responses.

Importantly, the functional preservation in both preparations does not invalidate the value of SEC purification. Higher purity preparations offer critical advantages for mechanistic studies, proteomic characterization, and therapeutic development where minimizing undefined variables is essential [31,32].

### 4.4 Inflammation-dependent dual effects on endothelial activation and barrier function

The most intriguing finding of this study is the paradoxical dual effect of HMEC-1-derived EVs on TNF-α pretreated endothelial cells: significant upregulation of VCAM-1 (2.0-fold for UC+SEC, 1.6-fold for UC Bulk) while simultaneously preserving VE-cadherin-mediated junction integrity (gap reduction from 25.0±2.5 μm² to 3.5–6.0 μm²). This observation is mechanistically unexpected because VCAM-1 upregulation typically correlates with endothelial dysfunction and increased permeability [36,37].

Our results appear to contradict the established paradigm that VCAM-1 expression and barrier disruption are coupled processes. However, several possibilities could reconcile this apparent paradox. First, VCAM-1 upregulation may represent a compensatory response to restore homeostasis rather than a marker of ongoing dysfunction. Wang et al. [38] demonstrated that LPS-stimulated endothelial EVs with high VCAM-1 expression activate NF-κB signaling in monocytes through integrin α4 interactions, suggesting VCAM-1+ EVs primarily mediate immune cell recruitment rather than direct barrier disruption. In our system, the VCAM-1 increase may signal immune surveillance activation while other EV components simultaneously stabilize junctions.

Second, the barrier-protective effect may be mediated by EV cargo distinct from inflammatory adhesion molecules. VE-cadherin junction integrity depends on the balance between disruptive forces (myosin light chain phosphorylation, actomyosin contraction) and stabilizing factors (cortactin localization, adherens junction assembly) [39,40]. Our proteomic analysis revealed enrichment of cytoskeletal scaffolding proteins (AHNAK, VIM, MYH9) and cell-cell adhesion components (desmoplakin, talin-1, filamin A) in EV preparations. These proteins could directly reinforce junctions even as inflammatory signaling increases VCAM-1 expression through parallel pathways.

Third, the strict inflammation-dependency we observed, where EVs had no detectable effect on healthy endothelial cells but potent effects on TNF-pretreated cells, suggests a threshold or permissive mechanism. The TNF-activated endothelium may upregulate specific receptors or alter membrane composition to enable EV uptake or signaling that does not occur in resting cells. This context-dependent responsiveness aligns with growing evidence that EV effects are highly dependent on recipient cell state [41,42,43]. Li et al. [44] demonstrated this principle dramatically in wound healing: fibroblast EVs from normal glucose conditions promoted angiogenesis while EVs from high-glucose conditions inhibited the same processes through opposing effects on GSK-3β/β-catenin signaling.

Our observation that VE-cadherin expression remained stable across all conditions while junction gaps decreased suggests EVs primarily affect junction assembly or stabilization rather than total VE-cadherin protein levels. Recent work showing EVs contain both soluble and membrane-anchored VE-cadherin, with membrane forms being functionally re-utilized by recipient cells [45], raises the possibility that EV-mediated VE-cadherin transfer could rapidly reinforce junctions independent of *de novo* protein synthesis.

### 4.5 Limitations and future directions

Several limitations warrant consideration. First, while we demonstrate functional effects of HMEC-1-derived EVs on endothelial activation and fibroblast migration, the molecular mechanisms underlying the paradoxical VCAM-1 upregulation and junction preservation remain undefined. Future studies should investigate whether specific EV proteins mediate these opposing effects, whether VCAM-1 blockade affects the barrier-protective phenotype, and which signaling pathways are activated in recipient cells.

Second, our functional assays used integrated cellular readouts (wound closure, junction gap measurement, VCAM-1 expression) that reflect multiple signaling pathways operating simultaneously. More mechanistic approaches, including receptor blockade experiments, pathway-specific inhibitors, and analysis of downstream signaling activation kinetics would provide clearer understanding of how EV cargo translates to cellular responses. The identification of 10 inflammatory response proteins in our proteomic analysis (adjusted p-value < 0.057) provides candidates for targeted functional validation.

Third, while we demonstrate that both UC Bulk and UC+SEC are functionally active in the specific assays tested, we did not perform comprehensive dose-response experiments that might reveal potency differences at limiting EV concentrations. The EV doses used (approximately 6–9.6×10□ nanoparticles per well, 60–96×10³ nanoparticles per cell) may saturate receptor-mediated responses, masking subtle differences in specific activity that would emerge at lower concentrations.

Finally, all functional experiments were performed *in vitro* using cell culture models. While HMEC-1 cells are well-validated for microvascular endothelial studies and primary human dermal microvascular endothelial cells (HDMECs) and human dermal fibroblasts (HDF) provided biological relevance, *in vivo* validation in preclinical models of inflammation or wound healing would strengthen translational significance. The inflammation-specific effects we observed suggest potential therapeutic applications in conditions characterized by endothelial activation, but efficacy and safety would require rigorous animal model testing.

## 5. CONCLUSIONS

We demonstrate that size exclusion chromatography following ultracentrifugation substantially improves EV purity through effective removal of serum proteins, extracellular matrix contaminants, and intracellular proteins while preserving core EV markers and functional activity. Both UC Bulk and UC+SEC preparations promote wound healing and exhibit unexpected dual effects on inflamed endothelial cells—simultaneously increasing VCAM-1 expression while preserving VE-cadherin junction integrity—effects that are strictly inflammation-dependent. Proteomic analysis reveals enrichment in wound healing, hemostasis, and angiogenesis pathways in both preparations despite compositional differences in specific proteins. These findings validate SEC as the preferred isolation method for high-purity EV preparations while demonstrating that ultracentrifugation-only preparations retain functional activity in specific biological assays. The inflammation-specific effects and paradoxical activation-barrier preservation phenotype warrant further mechanistic investigation to fully understand how EV cargo composition translates to complex cellular responses.

## Supporting information

Text References

## 6. ACKNOWLEDGMENTS

The authors thank the University of Virginia Electron Microscopy Core for technical support with cryogenic electron microscopy sample preparation and data acquisition, and the Georgetown University Medical Center Mass Spectrometry and Analytical Pharmacology Core for nanoUPLC-MS/MS analysis. This work was supported by the National Cancer Institute of the National Institutes of Health under award number U54 CA274499. The content is solely the responsibility of the authors and does not necessarily represent the official views of the National Institutes of Health.

