## Supplementary Figures for "HMEC-1 extracellular vesicles as regulators of endothelial cell activation under inflammation"

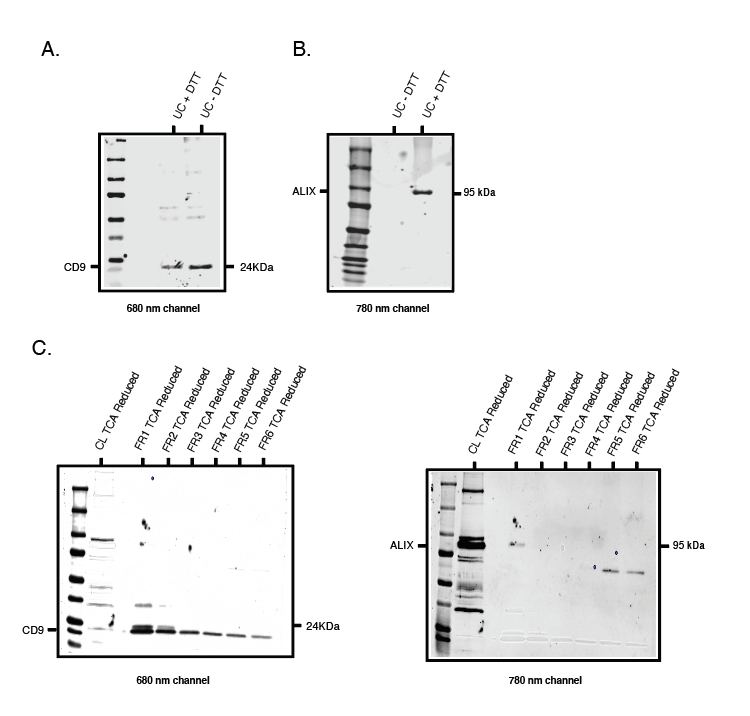

**Supplementary Figure 1. Full uncropped nitrocellulose membranes for EV marker characterization (corresponding to Figure 1B). (A) Full membrane of UC Bulk preparation probed for CD9 (24 kDa) detected on the 680 nm channel. UC Bulk samples were loaded under reducing (+DTT) and non-reducing (−DTT) conditions. (B) Full membrane of UC Bulk preparation probed for ALIX (95 kDa) detected on the 780 nm channel under reducing and non-reducing conditions. (C) Full membranes of HMEC-1 cell lysate (CL) and UC+SEC eluted fractions (FR1–FR6) co-stained for CD9 (680 nm channel, left) and ALIX (780 nm channel, right). All samples in (C) were concentrated by TCA/DOC precipitation prior to loading. Molecular weight markers are indicated.**

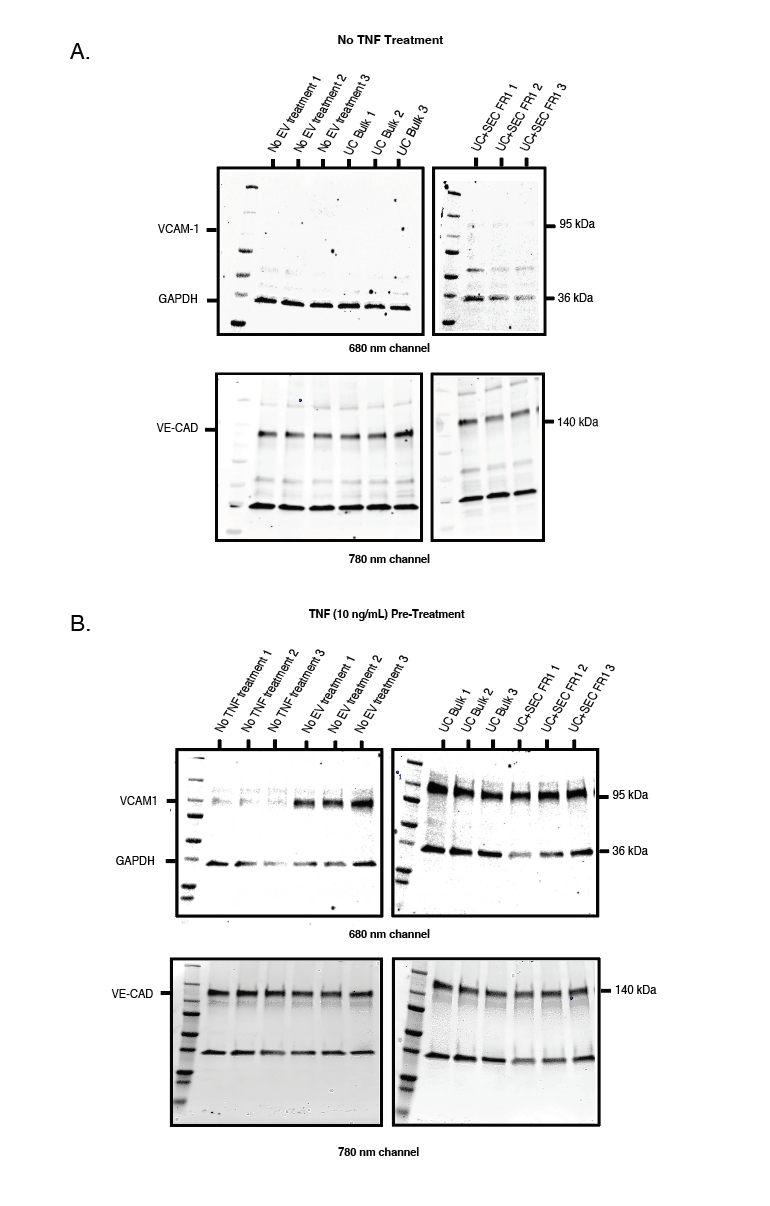

**Supplementary Figure 2. Full uncropped immunoblotting membranes for the endothelial activation assay (corresponding to Figure 3C and 3E). (A) Full membranes from healthy conditions (No TNF pre-treatment). Top: VCAM-1 (95 kDa) and GAPDH (36 kDa) detected on the 680 nm channel. Bottom: VE-cadherin (140 kDa) detected on the 780 nm channel. Lanes correspond to No EV treatment (n = 3), UC Bulk (n = 3), and UC+SEC FR1 (n = 3). (B) Full membranes from inflamed conditions (TNF 10 ng/mL pre-treatment). Top: VCAM-1 (95 kDa) and GAPDH (36 kDa) detected on the 680 nm channel. Bottom: VE-cadherin (140 kDa) detected on the 780 nm channel. Lanes correspond to No TNF treatment (n = 3), No EV treatment (n = 3), UC Bulk (n = 3), and UC+SEC FR1 (n = 3). Molecular weight markers are indicated. Imaging was performed on an Odyssey Infrared Imager (LI-COR Biosciences).**
