## Supplementary Tables for "HMEC-1 extracellular vesicles as regulators of endothelial cell activation under inflammation"

| **Name** | **Vendor** | **Vendor Stock** | **Reconstitution** | **For 10 mL** | **For 500 mL** | **Aliquot** | **Aliquot Conc.** | **Storage** |
| --- | --- | --- | --- | --- | --- | --- | --- | --- |
| rEGF | PrepoTech/Thermo | 100 µg | 100 µL sterile H₂O (1 mg/mL) then 1:10 in 0.1% BSA/H₂O for 1000 µL at 0.1 mg/mL (100 ng/µL) | At 10 ng/mL - 1 µL | At 10 ng/mL - 50 µL | 50 µL | 100 ng/µL | -80°C |
| bFGF | PrepoTech/Thermo | 50 µg | 50 µL sterile H₂O (1 mg/mL) then 1:10 in 0.1% BSA/H₂O for 500 µL at 0.1 mg/mL (100 ng/µL) | At 10 ng/mL - 1 µL | At 10 ng/mL - 50 µL | 50 µL | 100 ng/µL | -80°C |
| ANG1 | PrepoTech/Thermo | 20 µg | 20 µL sterile H₂O (1 mg/mL) then 1:20 in 0.1% BSA/H₂O for 400 µL at 0.05 mg/mL (50 ng/µL) | At 5 ng/mL - 1 µL | At 5 ng/mL - 50 µL | 50 µL | 50 ng/µL | -80°C |
| ANG2 | PrepoTech/Thermo | 5 µg | 50 µL sterile H₂O (0.1 mg/mL) then 1:10 in 0.1% BSA/H₂O for 500 µL at 0.01 mg/mL (10 ng/µL) | At 1 ng/mL - 1 µL | At 1 ng/mL - 50 µL | 50 µL | 10 ng/µL | -80°C |
| L-glutamine (GlutaMAX) | Gibco/Thermo | 200 mM | No need. | At 10 mM use 500 µL | At 10 mM use 25 mL | No need. | – | 4°C |
| Hydrocortisone | Sigma | 5 G | 40 mg in 10 mL of 95% ethanol. Filter 0.2 µm. | At 1 µg/mL use 2.5 µL | At 1 µg/mL use 125 µL | 125 µL | 4 mg/mL | -80°C |
| Ascorbic acid 2p (0.1 mM) | Sigma | 5–10 G | 1447 mg in 50 mL of basal media. Filter 0.2 µm. | At 0.1 mM use 10 µL | At 0.1 mM use 500 µL | 500 µL | 100 mM | 4°C |
| AlbuMAX | Sigma | 10 G | 1 g in 50 mL of basal media | At 0.5 mg/mL use 250 µL | At 0.5 mg/mL use 12.5 mL | 12500 µL | 20 mg/mL | -20°C |
| HEPES | Gibco/Thermo | 100X | No need. | At 5 mM use 50 µL | At 5 mM use 2500 µL | 5000 µL | 1M | 4°C |
| Dibutyryl-cAMP | Sigma | 100 mg | 100 mg in 10 mL of 0.1% BSA/H₂O. Filter 0.2 µm. | At 40.7 µM use 20 µL | At 40.7 µM use 1 mL | 1000 µL | 10 mg/mL | -20°C |
| IBMX | Sigma | 250 mg | 250 mg in 25 mL of sterile ethanol. | At 0.036 mM use 8 µL | At 0.036 mM use 368 µL | 368 µL | 10 mg/mL | -20°C |
| ITS (100x) | Gibco/Thermo | 100X | No need. | 50 µL for 0.5X | 2500 µL for 0.5X | 5000 µL | 0.5X | 4°C |
| Heparin | Sigma | 1000U–10000U | 500 mg in 10 mL | At 25 µg/mL use 5 µL | At 25 µg/mL use 250 µL | 250 µL | 50 mg/mL | 4°C |

*Supplementary Table 1: MCDB131 F15 supplementation reagents: vendors, reconstitution, and storage.*

| **Batch ID** | **Passage** | **Flask Model** | **Cell Seeding** | **Growth (h)** | **Collection (h)** | **Confluency (%)** | **Cell Count** | **pH** | **Viability (%)** | **NTA UC Bulk** | **NTA UC+SEC FR1** |
| --- | --- | --- | --- | --- | --- | --- | --- | --- | --- | --- | --- |
| ISO_1 | 8 | 1x T500 | 2.50E+06 | 72 | 48 | 95–100 | 2.50E+07 | 7.1 | 98% | 1.20E+10 | 7.00E+09 |
| ISO_2 | 7 | 1x T500 | 2.50E+06 | 72 | 48 | 95–100 | 2.50E+07 | 7.11 | 99% | 1.40E+10 | 9.00E+09 |
| ISO_3 | 10 | 3x T500 | 2.50E+06 | 72 | 48 | 95–100 | 2.30E+07 | 7.14 | 98% | 1.20E+10 (AVG) | – |
| ISO_4 | 10 | 3x T500 | 2.50E+06 | 72 | 48 | 95–100 | 2.50E+07 | 7.12 | 98% | 2.00E+10 (AVG) | 1.8E+10 (AVG) |
| ISO_5 | 12 | 8x T500 | 2.50E+06 | 72 | 48 | 95–100 | 2.00E+08 | 7.09 | 97% | 2.50E+11 | 7.00E+10 |
| ISO_6 | 14 | 8x T500 | 2.50E+06 | 72 | 48 | 95–100 | 2.00E+08 | 7.09 | 98% | 1.80E+11 | 5.00E+10 |
| ISO_7 | 6 | 8x T500 | 2.50E+06 | 72 | 48 | 95–100 | 1.80E+08 | 7.05 | 99% | 2.40E+11 | 6.54E+10 |
| ISO_8 | 8 | 8x T500 | 2.50E+06 | 72 | 48 | 95–100 | 1.80E+08 | 7.1 | 98% | 1.20E+11 | 5.40E+10 |

| **Batch ID** | **Batch Usage** |
| --- | --- |
| ISO_1 | Characterization: Cryo-EM, EV marker immunoblotting CD9, ALIX, TSG101 |
| ISO_2 | Characterization: EV marker immunoblotting CD81, HSP90. Micro-BCA quantification. |
| ISO_3 | Characterization: Mass spectrometry – UC Bulk |
| ISO_4 | Characterization: Mass spectrometry – UC Bulk+SEC FR1 |
| ISO_5 | Functionality: Pretreated TNF HDMEC model. |
| ISO_6 | Functionality: Fibroblast wound healing. Characterization: Micro-BCA quantification |
| ISO_7 | Functionality: Healthy (Non-treated) HDMEC model. |
| ISO_8 | Characterization: Micro-BCA quantification |

*Supplementary Table 2: HMEC-1 EV isolation batch records.*

**Supplementary Table 3**

| **Accession** | **Gene** | **Size (kDa)** | **UC Bulk Rep1** | **UC Bulk Rep2** | **UC Bulk Rep3** | **UC+SEC FR1 Rep1** | **UC+SEC FR1 Rep2** | **UC+SEC FR1 Rep3** |
| --- | --- | --- | --- | --- | --- | --- | --- | --- |
| B5ME19 | EIF3CL | 105.4 | 0 | 0 | 0 | 3 | 0 | 7 |
| O00151 | PDLIM1 | 36 | 0 | 0 | 4 | 0 | 0 | 3 |
| O00159 | MYO1C | 121.6 | 0 | 3 | 8 | 0 | 6 | 25 |
| O00161 | SNAP23 | 23.3 | 0 | 0 | 0 | 3 | 0 | 4 |
| O00186 | STXBP3 | 67.7 | 0 | 0 | 0 | 0 | 0 | 7 |
| O00203 | AP3B1 | 121.2 | 0 | 0 | 0 | 0 | 0 | 3 |
| O00231 | PSMD11 | 47.4 | 0 | 0 | 0 | 0 | 0 | 7 |
| O00232 | PSMD12 | 52.9 | 0 | 0 | 0 | 0 | 0 | 4 |
| O00299 | CLIC1 | 26.9 | 0 | 5 | 7 | 5 | 4 | 10 |
| O00391 | QSOX1 | 82.5 | 0 | 3 | 3 | 0 | 0 | 5 |
| O00468 | AGRN | 217.2 | 0 | 0 | 4 | 0 | 0 | 20 |
| O00487 | PSMD14 | 34.6 | 0 | 0 | 0 | 3 | 0 | 4 |
| O00560 | SDCBP | 32.4 | 5 | 5 | 7 | 6 | 5 | 14 |
| O00571 | DDX3X | 73.2 | 0 | 0 | 0 | 0 | 0 | 5 |
| O00592 | PODXL | 58.6 | 0 | 0 | 0 | 0 | 0 | 3 |
| O00622 | CCN1 | 42 | 5 | 4 | 7 | 0 | 0 | 9 |
| O14672 | ADAM10 | 84.1 | 0 | 0 | 0 | 0 | 0 | 12 |
| O14745 | SLC9A3R1 | 38.8 | 0 | 0 | 0 | 0 | 0 | 3 |
| O14786 | NRP1 | 103.1 | 0 | 0 | 5 | 0 | 0 | 4 |
| O14828 | SCAMP3 | 38.3 | 0 | 0 | 0 | 0 | 0 | 3 |
| O14950 | MYL12B | 19.8 | 0 | 0 | 0 | 0 | 0 | 5 |
| O15143 | ARPC1B | 40.9 | 0 | 0 | 0 | 0 | 0 | 6 |
| O15144 | ARPC2 | 34.3 | 0 | 0 | 0 | 0 | 0 | 3 |
| O15145 | ARPC3 | 20.5 | 0 | 0 | 0 | 0 | 0 | 4 |
| O15162 | PLSCR1 | 35 | 0 | 0 | 0 | 3 | 4 | 6 |
| O15230 | LAMA5 | 399.5 | 0 | 0 | 5 | 0 | 0 | 26 |
| O15260 | SURF4 | 30.4 | 0 | 0 | 0 | 0 | 0 | 3 |
| O15294 | OGT | 116.9 | 0 | 9 | 0 | 0 | 0 | 0 |
| O15427 | SLC16A3 | 49.4 | 0 | 0 | 0 | 4 | 0 | 4 |
| O43143 | DHX15 | 90.9 | 0 | 0 | 0 | 0 | 0 | 6 |
| O43175 | PHGDH | 56.6 | 0 | 0 | 0 | 0 | 0 | 6 |
| O43242 | PSMD3 | 60.9 | 0 | 0 | 0 | 3 | 0 | 9 |
| O43390 | HNRNPR | 70.9 | 0 | 0 | 0 | 0 | 0 | 6 |
| O43488 | AKR7A2 | 39.6 | 0 | 0 | 0 | 0 | 0 | 3 |
| O43491 | EPB41L2 | 112.5 | 0 | 0 | 0 | 0 | 0 | 3 |
| O43633 | CHMP2A | 25.1 | 0 | 0 | 0 | 0 | 0 | 3 |
| O43657 | TSPAN6 | 27.5 | 0 | 0 | 0 | 0 | 0 | 3 |
| O43684 | BUB3 | 37.1 | 0 | 0 | 0 | 0 | 0 | 8 |
| O43707 | ACTN4 | 104.8 | 0 | 3 | 4 | 3 | 0 | 10 |
| O43747 | AP1G1 | 91.3 | 0 | 0 | 0 | 0 | 0 | 5 |
| O43776 | NARS1 | 62.9 | 3 | 0 | 0 | 0 | 0 | 9 |
| O43795 | MYO1B | 131.9 | 0 | 0 | 0 | 0 | 0 | 5 |
| O43854 | EDIL3 | 53.7 | 10 | 11 | 14 | 0 | 0 | 16 |
| O43865 | AHCYL1 | 58.9 | 0 | 0 | 0 | 0 | 0 | 3 |
| O60264 | SMARCA5 | 121.8 | 0 | 0 | 0 | 0 | 0 | 3 |
| O60487 | MPZL2 | 24.5 | 0 | 0 | 0 | 0 | 0 | 3 |
| O60488 | ACSL4 | 79.1 | 0 | 0 | 0 | 0 | 0 | 5 |
| O60568 | PLOD3 | 84.7 | 0 | 0 | 3 | 0 | 0 | 0 |
| O60701 | UGDH | 55 | 0 | 0 | 0 | 0 | 0 | 7 |
| O60716 | CTNND1 | 108.1 | 3 | 0 | 3 | 0 | 0 | 6 |
| O60884 | DNAJA2 | 45.7 | 0 | 0 | 0 | 0 | 0 | 3 |
| O75044 | SRGAP2 | 120.8 | 0 | 0 | 0 | 0 | 0 | 3 |
| O75051 | PLXNA2 | 211 | 0 | 0 | 3 | 0 | 0 | 10 |
| O75083 | WDR1 | 66.2 | 3 | 5 | 5 | 0 | 0 | 8 |
| O75131 | CPNE3 | 60.1 | 0 | 0 | 6 | 8 | 5 | 16 |
| O75173 | ADAMTS4 | 90.1 | 0 | 0 | 3 | 0 | 0 | 4 |
| O75340 | PDCD6 | 21.9 | 0 | 0 | 0 | 7 | 5 | 8 |
| O75351 | VPS4B | 49.3 | 0 | 0 | 0 | 0 | 0 | 5 |
| O75367 | MACROH2A1 | 39.2 | 0 | 0 | 0 | 3 | 0 | 0 |
| O75369 | FLNB | 278 | 6 | 3 | 10 | 10 | 5 | 20 |
| O75390 | CS | 51.7 | 0 | 0 | 0 | 0 | 0 | 4 |
| O75533 | SF3B1 | 145.7 | 0 | 0 | 0 | 0 | 0 | 4 |
| O75643 | SNRNP200 | 244.4 | 0 | 0 | 0 | 0 | 0 | 6 |
| O75695 | RP2 | 39.6 | 0 | 0 | 0 | 0 | 0 | 4 |
| O75923 | DYSF | 237.1 | 0 | 0 | 0 | 0 | 0 | 17 |
| O75954 | TSPAN9 | 26.8 | 0 | 0 | 0 | 0 | 0 | 3 |
| O75955 | FLOT1 | 47.3 | 0 | 0 | 0 | 0 | 0 | 9 |
| O76021 | RSL1D1 | 54.9 | 0 | 0 | 0 | 0 | 0 | 8 |
| O94760 | DDAH1 | 31.1 | 0 | 0 | 0 | 0 | 0 | 3 |
| O94776 | MTA2 | 75 | 0 | 0 | 0 | 0 | 0 | 5 |
| O94813 | SLIT2 | 169.8 | 9 | 4 | 15 | 0 | 0 | 14 |
| O94979 | SEC31A | 132.9 | 0 | 0 | 0 | 0 | 0 | 3 |
| O95084 | PRSS23 | 43 | 0 | 0 | 5 | 4 | 0 | 4 |
| O95297 | MPZL1 | 29.1 | 0 | 0 | 0 | 0 | 0 | 3 |
| O95445 | APOM | 21.2 | 3 | 3 | 3 | 0 | 0 | 0 |
| O95782 | AP2A1 | 107.5 | 0 | 0 | 0 | 0 | 0 | 4 |
| O95810 | CAVIN2 | 47.1 | 0 | 0 | 0 | 0 | 0 | 3 |
| O95819 | MAP4K4 | 142 | 0 | 0 | 3 | 0 | 0 | 6 |
| P00338 | LDHA | 36.7 | 0 | 4 | 3 | 4 | 0 | 7 |
| P00352 | ALDH1A1 | 54.8 | 3 | 4 | 5 | 4 | 0 | 8 |
| P00450 | CP | 122.1 | 0 | 0 | 3 | 0 | 0 | 0 |
| P00491 | PNP | 32.1 | 0 | 3 | 4 | 4 | 0 | 5 |
| P00558 | PGK1 | 44.6 | 5 | 5 | 8 | 6 | 4 | 12 |
| P00568 | AK1 | 21.6 | 0 | 0 | 0 | 0 | 0 | 3 |
| P00734 | F2 | 70 | 5 | 6 | 7 | 5 | 4 | 7 |
| P00738 | HP | 45.2 | 0 | 0 | 0 | 0 | 0 | 3 |
| P00750 | PLAT | 62.9 | 3 | 7 | 9 | 4 | 0 | 9 |
| P01008 | SERPINC1 | 52.6 | 3 | 3 | 4 | 3 | 0 | 4 |
| P01023 | A2M | 163.2 | 0 | 0 | 3 | 0 | 3 | 5 |
| P01024 | C3 | 187 | 12 | 12 | 12 | 0 | 0 | 9 |
| P01031 | C5 | 188.2 | 3 | 0 | 4 | 0 | 0 | 0 |
| P01111 | NRAS | 21.2 | 0 | 0 | 3 | 0 | 0 | 3 |
| P01857 | IGHG1 | 36.1 | 0 | 0 | 0 | 0 | 0 | 3 |
| P01889 | HLA-B | 40.4 | 0 | 0 | 3 | 0 | 0 | 0 |
| P02452 | COL1A1 | 138.8 | 10 | 4 | 13 | 0 | 0 | 4 |
| P02545 | LMNA | 74.1 | 4 | 6 | 6 | 9 | 4 | 17 |
| P02649 | APOE | 36.1 | 3 | 3 | 5 | 4 | 0 | 5 |
| P02748 | C9 | 63.1 | 0 | 0 | 0 | 0 | 0 | 3 |
| P02751 | FN1 | 272.2 | 69 | 68 | 80 | 38 | 43 | 71 |
| P02765 | AHSG | 39.3 | 4 | 3 | 5 | 0 | 0 | 0 |
| P02768 | ALB | 69.3 | 8 | 8 | 9 | 7 | 7 | 16 |
| P02771 | AFP | 68.6 | 0 | 4 | 0 | 0 | 0 | 0 |
| P02774 | GC | 52.9 | 4 | 3 | 4 | 0 | 0 | 0 |
| P02786 | TFRC | 84.8 | 0 | 0 | 0 | 0 | 0 | 8 |
| P02787 | TF | 77 | 31 | 36 | 34 | 0 | 0 | 16 |
| P02788 | LTF | 78.1 | 3 | 0 | 3 | 0 | 0 | 0 |
| P02790 | HPX | 51.6 | 6 | 3 | 6 | 7 | 6 | 15 |
| P02794 | FTH1 | 21.2 | 0 | 0 | 0 | 0 | 0 | 3 |
| P03956 | MMP1 | 54 | 3 | 3 | 4 | 0 | 0 | 0 |
| P04004 | VTN | 54.3 | 0 | 0 | 0 | 0 | 0 | 3 |
| P04040 | CAT | 59.7 | 7 | 0 | 0 | 3 | 0 | 0 |
| P04075 | ALDOA | 39.4 | 3 | 6 | 8 | 7 | 0 | 15 |
| P04083 | ANXA1 | 38.7 | 11 | 16 | 15 | 18 | 16 | 25 |
| P04114 | APOB | 515.3 | 15 | 15 | 15 | 4 | 3 | 9 |
| P04275 | VWF | 309.1 | 0 | 0 | 4 | 0 | 0 | 0 |
| P04406 | GAPDH | 36 | 11 | 12 | 14 | 15 | 10 | 23 |
| P04439 | HLA-A | 40.8 | 0 | 0 | 4 | 3 | 0 | 7 |
| P04792 | HSPB1 | 22.8 | 3 | 3 | 5 | 3 | 0 | 3 |
| P04899 | GNAI2 | 40.4 | 8 | 7 | 8 | 3 | 0 | 9 |
| P05023 | ATP1A1 | 112.8 | 9 | 9 | 13 | 13 | 10 | 26 |
| P05026 | ATP1B1 | 35 | 0 | 0 | 0 | 0 | 0 | 4 |
| P05089 | ARG1 | 34.7 | 4 | 0 | 0 | 7 | 0 | 5 |
| P05106 | ITGB3 | 87 | 0 | 0 | 0 | 0 | 0 | 9 |
| P05109 | S100A8 | 10.8 | 0 | 0 | 0 | 0 | 0 | 3 |
| P05121 | SERPINE1 | 45 | 10 | 11 | 11 | 4 | 0 | 12 |
| P05198 | EIF2S1 | 36.1 | 0 | 0 | 0 | 0 | 0 | 3 |
| P05362 | ICAM1 | 57.8 | 0 | 0 | 0 | 3 | 0 | 3 |
| P05387 | RPLP2 | 11.7 | 0 | 0 | 0 | 4 | 0 | 3 |
| P05388 | RPLP0 | 34.3 | 5 | 4 | 6 | 3 | 3 | 3 |
| P05452 | CLEC3B | 22.5 | 0 | 3 | 3 | 0 | 0 | 0 |
| P05556 | ITGB1 | 88.4 | 11 | 8 | 12 | 12 | 11 | 22 |
| P06396 | GSN | 85.6 | 6 | 7 | 8 | 0 | 0 | 5 |
| P06493 | CDK1 | 34.1 | 0 | 0 | 0 | 0 | 0 | 6 |
| P06702 | S100A9 | 13.2 | 0 | 0 | 0 | 3 | 0 | 0 |
| P06733 | ENO1 | 47.1 | 11 | 12 | 16 | 15 | 6 | 23 |
| P06737 | PYGL | 97.1 | 0 | 0 | 0 | 0 | 0 | 7 |
| P06744 | GPI | 63.1 | 0 | 0 | 3 | 0 | 0 | 7 |
| P06748 | NPM1 | 32.6 | 0 | 0 | 0 | 4 | 0 | 3 |
| P06753 | TPM3 | 32.9 | 0 | 0 | 0 | 0 | 0 | 3 |
| P06756 | ITGAV | 116 | 3 | 0 | 6 | 0 | 0 | 18 |
| P07195 | LDHB | 36.6 | 4 | 3 | 7 | 6 | 0 | 9 |
| P07339 | CTSD | 44.5 | 4 | 0 | 0 | 0 | 0 | 0 |
| P07355 | ANXA2 | 38.6 | 21 | 28 | 25 | 24 | 21 | 37 |
| P07384 | CAPN1 | 81.8 | 0 | 0 | 0 | 0 | 0 | 5 |
| P07437 | TUBB | 49.6 | 4 | 4 | 4 | 3 | 4 | 4 |
| P07737 | PFN1 | 15 | 0 | 4 | 3 | 5 | 3 | 9 |
| P07814 | EPRS1 | 170.5 | 0 | 0 | 4 | 0 | 0 | 16 |
| P07900 | HSP90AA1 | 84.6 | 9 | 8 | 9 | 7 | 6 | 11 |
| P07910 | HNRNPC | 33.7 | 0 | 0 | 0 | 4 | 0 | 6 |
| P07942 | LAMB1 | 197.9 | 7 | 4 | 12 | 0 | 0 | 14 |
| P07947 | YES1 | 60.8 | 4 | 0 | 0 | 0 | 0 | 6 |
| P07996 | THBS1 | 129.3 | 36 | 34 | 39 | 12 | 10 | 40 |
| P08133 | ANXA6 | 75.8 | 6 | 7 | 12 | 11 | 9 | 34 |
| P08134 | RHOC | 22 | 0 | 4 | 0 | 0 | 0 | 0 |
| P08195 | SLC3A2 | 68 | 0 | 0 | 0 | 4 | 4 | 10 |
| P08237 | PFKM | 85.1 | 0 | 0 | 0 | 0 | 0 | 6 |
| P08238 | HSP90AB1 | 83.2 | 10 | 10 | 9 | 9 | 6 | 15 |
| P08253 | MMP2 | 73.8 | 3 | 4 | 5 | 0 | 0 | 0 |
| P08572 | COL4A2 | 167.4 | 0 | 0 | 0 | 0 | 0 | 5 |
| P08621 | SNRNP70 | 51.5 | 0 | 0 | 0 | 0 | 0 | 3 |
| P08648 | ITGA5 | 114.5 | 8 | 4 | 8 | 6 | 3 | 11 |
| P08670 | VIM | 53.6 | 5 | 6 | 12 | 17 | 15 | 18 |
| P08708 | RPS17 | 15.5 | 3 | 3 | 3 | 0 | 0 | 7 |
| P08758 | ANXA5 | 35.9 | 8 | 13 | 15 | 12 | 9 | 20 |
| P08779 | KRT16 | 51.2 | 0 | 0 | 0 | 0 | 0 | 9 |
| P08865 | RPSA | 32.8 | 4 | 6 | 7 | 8 | 3 | 9 |
| P09038 | FGF2 | 30.8 | 0 | 0 | 3 | 0 | 0 | 4 |
| P09211 | GSTP1 | 23.3 | 0 | 0 | 0 | 0 | 0 | 5 |
| P09382 | LGALS1 | 14.7 | 0 | 0 | 3 | 6 | 3 | 7 |
| P09486 | SPARC | 34.6 | 0 | 0 | 4 | 0 | 0 | 0 |
| P09496 | CLTA | 27.1 | 0 | 0 | 0 | 0 | 0 | 5 |
| P09525 | ANXA4 | 35.9 | 0 | 0 | 0 | 7 | 5 | 14 |
| P09543 | CNP | 47.5 | 3 | 0 | 3 | 3 | 0 | 4 |
| P09874 | PARP1 | 113 | 0 | 0 | 0 | 0 | 0 | 5 |
| P0C0L4 | C4A | 192.7 | 8 | 6 | 5 | 0 | 0 | 4 |
| P0DMV9 | HSPA1B | 70 | 5 | 7 | 8 | 9 | 5 | 17 |
| P0DP25 | CALM3 | 16.8 | 0 | 0 | 0 | 5 | 7 | 5 |
| P0DUB6 | AMY1A | 57.7 | 0 | 0 | 3 | 0 | 0 | 0 |
| P10301 | RRAS | 23.5 | 0 | 0 | 0 | 0 | 0 | 4 |
| P10646 | TFPI | 35 | 0 | 0 | 3 | 0 | 0 | 3 |
| P10768 | ESD | 31.4 | 0 | 0 | 0 | 0 | 0 | 3 |
| P10915 | HAPLN1 | 40.1 | 0 | 0 | 0 | 0 | 0 | 3 |
| P11021 | HSPA5 | 72.3 | 3 | 0 | 3 | 0 | 0 | 9 |
| P11047 | LAMC1 | 177.5 | 8 | 3 | 14 | 0 | 0 | 15 |
| P11142 | HSPA8 | 70.9 | 15 | 18 | 19 | 17 | 15 | 25 |
| P11166 | SLC2A1 | 54 | 0 | 0 | 0 | 0 | 0 | 4 |
| P11216 | PYGB | 96.6 | 0 | 0 | 0 | 5 | 0 | 11 |
| P11233 | RALA | 23.6 | 0 | 0 | 3 | 0 | 0 | 3 |
| P11387 | TOP1 | 90.7 | 0 | 0 | 0 | 4 | 0 | 8 |
| P11388 | TOP2A | 174.3 | 0 | 0 | 0 | 7 | 0 | 13 |
| P11413 | G6PD | 59.2 | 0 | 0 | 0 | 0 | 0 | 3 |
| P11586 | MTHFD1 | 101.5 | 3 | 0 | 0 | 5 | 0 | 11 |
| P11940 | PABPC1 | 70.6 | 0 | 0 | 0 | 0 | 0 | 9 |
| P12004 | PCNA | 28.8 | 0 | 0 | 0 | 0 | 0 | 4 |
| P12109 | COL6A1 | 108.5 | 4 | 3 | 6 | 0 | 0 | 4 |
| P12111 | COL6A3 | 343.5 | 0 | 0 | 0 | 0 | 0 | 9 |
| P12259 | F5 | 251.5 | 3 | 0 | 3 | 5 | 0 | 5 |
| P12268 | IMPDH2 | 55.8 | 0 | 0 | 3 | 0 | 0 | 0 |
| P12273 | PIP | 16.6 | 0 | 0 | 0 | 0 | 0 | 4 |
| P12429 | ANXA3 | 36.4 | 0 | 3 | 5 | 5 | 4 | 13 |
| P12814 | ACTN1 | 103 | 0 | 0 | 7 | 3 | 7 | 8 |
| P12956 | XRCC6 | 69.8 | 0 | 0 | 0 | 4 | 0 | 9 |
| P13010 | XRCC5 | 82.7 | 0 | 0 | 0 | 0 | 0 | 5 |
| P13489 | RNH1 | 49.9 | 0 | 0 | 0 | 0 | 0 | 5 |
| P13612 | ITGA4 | 114.8 | 0 | 0 | 0 | 0 | 0 | 3 |
| P13639 | EEF2 | 95.3 | 13 | 11 | 19 | 20 | 6 | 36 |
| P13797 | PLS3 | 70.8 | 4 | 5 | 5 | 0 | 0 | 7 |
| P13987 | CD59 | 14.2 | 0 | 0 | 0 | 0 | 3 | 5 |
| P14324 | FDPS | 48.2 | 0 | 0 | 0 | 0 | 0 | 5 |
| P14543 | NID1 | 136.3 | 5 | 9 | 10 | 0 | 0 | 13 |
| P14618 | PKM | 57.9 | 13 | 16 | 18 | 21 | 15 | 33 |
| P14625 | HSP90B1 | 92.4 | 0 | 0 | 0 | 0 | 0 | 3 |
| P14866 | HNRNPL | 64.1 | 0 | 0 | 0 | 0 | 0 | 4 |
| P14868 | DARS1 | 57.1 | 0 | 0 | 3 | 4 | 0 | 11 |
| P14923 | JUP | 81.7 | 11 | 0 | 11 | 15 | 3 | 18 |
| P15121 | AKR1B1 | 35.8 | 0 | 0 | 0 | 3 | 0 | 4 |
| P15144 | ANPEP | 109.5 | 3 | 3 | 8 | 13 | 12 | 26 |
| P15311 | EZR | 69.4 | 0 | 0 | 0 | 8 | 6 | 13 |
| P15559 | NQO1 | 30.8 | 0 | 0 | 0 | 0 | 0 | 3 |
| P15880 | RPS2 | 31.3 | 0 | 0 | 0 | 10 | 4 | 12 |
| P15924 | DSP | 331.6 | 29 | 0 | 8 | 27 | 6 | 24 |
| P16070 | CD44 | 81.5 | 0 | 0 | 3 | 5 | 0 | 5 |
| P16152 | CBR1 | 30.4 | 0 | 0 | 0 | 3 | 0 | 6 |
| P16284 | PECAM1 | 82.5 | 0 | 0 | 4 | 3 | 0 | 9 |
| P16949 | STMN1 | 17.3 | 0 | 0 | 0 | 3 | 0 | 0 |
| P17252 | PRKCA | 76.7 | 0 | 0 | 0 | 0 | 0 | 3 |
| P17301 | ITGA2 | 129.2 | 6 | 3 | 11 | 0 | 0 | 10 |
| P17302 | GJA1 | 43 | 0 | 0 | 0 | 0 | 0 | 4 |
| P17612 | PRKACA | 40.6 | 0 | 0 | 0 | 0 | 0 | 3 |
| P17655 | CAPN2 | 79.9 | 0 | 0 | 0 | 0 | 0 | 3 |
| P17813 | ENG | 70.5 | 0 | 3 | 4 | 3 | 0 | 9 |
| P17844 | DDX5 | 69.1 | 0 | 0 | 0 | 0 | 0 | 6 |
| P17858 | PFKL | 85 | 0 | 0 | 0 | 0 | 0 | 6 |
| P17931 | LGALS3 | 26.1 | 0 | 0 | 0 | 0 | 0 | 6 |
| P17980 | PSMC3 | 49.2 | 0 | 0 | 0 | 0 | 0 | 6 |
| P17987 | TCP1 | 60.3 | 0 | 0 | 8 | 7 | 3 | 8 |
| P18077 | RPL35A | 12.5 | 0 | 0 | 0 | 3 | 0 | 0 |
| P18085 | ARF4 | 20.5 | 0 | 0 | 0 | 0 | 0 | 3 |
| P18124 | RPL7 | 29.2 | 4 | 4 | 6 | 14 | 6 | 17 |
| P18206 | VCL | 123.7 | 12 | 12 | 13 | 9 | 8 | 23 |
| P18621 | RPL17 | 21.4 | 0 | 0 | 3 | 4 | 0 | 7 |
| P18669 | PGAM1 | 28.8 | 3 | 4 | 3 | 3 | 0 | 5 |
| P19338 | NCL | 76.6 | 0 | 0 | 0 | 6 | 0 | 10 |
| P19823 | ITIH2 | 106.4 | 5 | 6 | 6 | 0 | 0 | 5 |
| P20073 | ANXA7 | 52.7 | 3 | 0 | 3 | 6 | 3 | 13 |
| P20338 | RAB4A | 24.4 | 0 | 0 | 0 | 0 | 0 | 3 |
| P20339 | RAB5A | 23.6 | 0 | 0 | 0 | 0 | 0 | 3 |
| P20340 | RAB6A | 23.6 | 0 | 0 | 0 | 0 | 0 | 3 |
| P20591 | MX1 | 75.5 | 0 | 0 | 0 | 4 | 0 | 0 |
| P20618 | PSMB1 | 26.5 | 0 | 0 | 3 | 0 | 0 | 0 |
| P20774 | OGN | 33.9 | 3 | 0 | 0 | 0 | 0 | 0 |
| P20908 | COL5A1 | 183.4 | 0 | 0 | 3 | 0 | 0 | 0 |
| P21333 | FLNA | 280.6 | 26 | 15 | 36 | 28 | 14 | 50 |
| P21399 | ACO1 | 98.3 | 0 | 0 | 0 | 0 | 0 | 10 |
| P21589 | NT5E | 63.3 | 9 | 7 | 12 | 12 | 11 | 22 |
| P21926 | CD9 | 25.4 | 3 | 4 | 4 | 5 | 5 | 9 |
| P21980 | TGM2 | 77.3 | 8 | 8 | 11 | 5 | 3 | 10 |
| P22102 | GART | 107.7 | 0 | 0 | 0 | 0 | 0 | 3 |
| P22105 | TNXB | 458.1 | 5 | 0 | 3 | 0 | 0 | 0 |
| P22234 | PAICS | 47 | 0 | 0 | 3 | 4 | 0 | 6 |
| P22314 | UBA1 | 117.8 | 6 | 3 | 7 | 5 | 0 | 12 |
| P22626 | HNRNPA2B1 | 37.4 | 0 | 0 | 3 | 4 | 0 | 4 |
| P23142 | FBLN1 | 77.2 | 7 | 5 | 8 | 0 | 0 | 0 |
| P23229 | ITGA6 | 126.5 | 0 | 0 | 3 | 0 | 0 | 9 |
| P23246 | SFPQ | 76.1 | 0 | 0 | 3 | 0 | 0 | 4 |
| P23396 | RPS3 | 26.7 | 6 | 3 | 9 | 12 | 0 | 15 |
| P23526 | AHCY | 47.7 | 7 | 3 | 7 | 0 | 0 | 9 |
| P23528 | CFL1 | 18.5 | 0 | 4 | 4 | 7 | 5 | 6 |
| P23634 | ATP2B4 | 137.8 | 0 | 0 | 0 | 0 | 0 | 11 |
| P24844 | MYL9 | 19.8 | 0 | 0 | 0 | 3 | 0 | 0 |
| P25311 | AZGP1 | 34.2 | 0 | 0 | 0 | 0 | 0 | 3 |
| P25398 | RPS12 | 14.5 | 0 | 0 | 0 | 0 | 0 | 4 |
| P26006 | ITGA3 | 116.5 | 6 | 4 | 8 | 0 | 3 | 11 |
| P26022 | PTX3 | 41.9 | 5 | 7 | 6 | 5 | 6 | 11 |
| P26038 | MSN | 67.8 | 12 | 16 | 16 | 17 | 9 | 26 |
| P26368 | U2AF2 | 53.5 | 0 | 0 | 0 | 0 | 0 | 3 |
| P26373 | RPL13 | 24.2 | 0 | 3 | 3 | 5 | 0 | 8 |
| P26639 | TARS1 | 83.4 | 0 | 0 | 0 | 0 | 0 | 3 |
| P26641 | EEF1G | 50.1 | 0 | 0 | 5 | 5 | 0 | 9 |
| P27105 | STOM | 31.7 | 5 | 4 | 7 | 9 | 7 | 15 |
| P27348 | YWHAQ | 27.7 | 3 | 0 | 0 | 4 | 3 | 4 |
| P27487 | DPP4 | 88.2 | 0 | 0 | 3 | 0 | 0 | 10 |
| P27635 | RPL10 | 24.6 | 0 | 0 | 3 | 4 | 0 | 8 |
| P27701 | CD82 | 29.6 | 0 | 0 | 0 | 0 | 0 | 4 |
| P28482 | MAPK1 | 41.4 | 0 | 0 | 0 | 0 | 0 | 4 |
| P29144 | TPP2 | 138.3 | 12 | 11 | 20 | 0 | 0 | 3 |
| P29279 | CCN2 | 38.1 | 7 | 9 | 7 | 0 | 0 | 6 |
| P29317 | EPHA2 | 108.2 | 3 | 0 | 7 | 4 | 3 | 15 |
| P29323 | EPHB2 | 117.4 | 0 | 0 | 0 | 0 | 0 | 3 |
| P29401 | TKT | 67.8 | 5 | 8 | 10 | 5 | 0 | 11 |
| P29692 | EEF1D | 31.1 | 0 | 0 | 0 | 4 | 0 | 4 |
| P29966 | MARCKS | 31.5 | 0 | 0 | 0 | 5 | 4 | 4 |
| P29992 | GNA11 | 42.1 | 0 | 0 | 0 | 0 | 3 | 3 |
| P30041 | PRDX6 | 25 | 0 | 0 | 0 | 0 | 0 | 5 |
| P30050 | RPL12 | 17.8 | 6 | 6 | 5 | 4 | 0 | 4 |
| P30086 | PEBP1 | 21 | 0 | 0 | 0 | 0 | 0 | 4 |
| P30101 | PDIA3 | 56.7 | 0 | 0 | 0 | 4 | 0 | 9 |
| P30153 | PPP2R1A | 65.3 | 0 | 0 | 3 | 3 | 0 | 6 |
| P30626 | SRI | 21.7 | 0 | 0 | 0 | 4 | 3 | 5 |
| P31151 | S100A7 | 11.5 | 0 | 0 | 0 | 0 | 3 | 3 |
| P31153 | MAT2A | 43.6 | 0 | 0 | 0 | 0 | 0 | 3 |
| P31689 | DNAJA1 | 44.8 | 0 | 0 | 4 | 3 | 4 | 8 |
| P31939 | ATIC | 64.6 | 0 | 0 | 0 | 0 | 0 | 6 |
| P31943 | HNRNPH1 | 49.2 | 0 | 0 | 0 | 0 | 0 | 3 |
| P31944 | CASP14 | 27.7 | 3 | 0 | 0 | 8 | 0 | 3 |
| P31946 | YWHAB | 28.1 | 0 | 0 | 0 | 4 | 3 | 4 |
| P31948 | STIP1 | 62.6 | 0 | 0 | 0 | 0 | 0 | 5 |
| P31949 | S100A11 | 11.7 | 0 | 0 | 0 | 0 | 3 | 3 |
| P32119 | PRDX2 | 21.9 | 3 | 0 | 0 | 5 | 0 | 5 |
| P32969 | RPL9P9 | 21.9 | 0 | 0 | 0 | 4 | 0 | 5 |
| P33176 | KIF5B | 109.6 | 0 | 0 | 0 | 3 | 0 | 9 |
| P34897 | SHMT2 | 56 | 0 | 0 | 0 | 0 | 0 | 5 |
| P34932 | HSPA4 | 94.3 | 0 | 0 | 0 | 0 | 0 | 4 |
| P35221 | CTNNA1 | 100 | 3 | 0 | 11 | 4 | 3 | 18 |
| P35222 | CTNNB1 | 85.4 | 3 | 3 | 0 | 0 | 0 | 4 |
| P35241 | RDX | 68.5 | 0 | 0 | 0 | 3 | 0 | 10 |
| P35268 | RPL22 | 14.8 | 0 | 0 | 0 | 0 | 0 | 3 |
| P35443 | THBS4 | 105.8 | 0 | 0 | 3 | 0 | 0 | 0 |
| P35579 | MYH9 | 226.4 | 29 | 18 | 36 | 52 | 26 | 86 |
| P35580 | MYH10 | 228.9 | 0 | 0 | 0 | 0 | 0 | 4 |
| P35606 | COPB2 | 102.4 | 0 | 0 | 0 | 0 | 0 | 6 |
| P35613 | BSG | 42.2 | 5 | 4 | 5 | 4 | 6 | 8 |
| P35659 | DEK | 42.6 | 0 | 0 | 0 | 0 | 0 | 4 |
| P35998 | PSMC2 | 48.6 | 0 | 0 | 3 | 0 | 0 | 10 |
| P36578 | RPL4 | 47.7 | 6 | 7 | 11 | 13 | 7 | 21 |
| P36955 | SERPINF1 | 46.3 | 4 | 3 | 3 | 0 | 0 | 0 |
| P37802 | TAGLN2 | 22.4 | 0 | 0 | 5 | 3 | 0 | 9 |
| P37837 | TALDO1 | 37.5 | 0 | 0 | 0 | 0 | 0 | 5 |
| P38919 | EIF4A3 | 46.8 | 0 | 0 | 0 | 3 | 0 | 0 |
| P39019 | RPS19 | 16.1 | 3 | 3 | 6 | 3 | 0 | 4 |
| P39023 | RPL3 | 46.1 | 6 | 10 | 9 | 8 | 3 | 12 |
| P39060 | COL18A1 | 178.1 | 9 | 11 | 9 | 0 | 0 | 0 |
| P40227 | CCT6A | 58 | 4 | 0 | 6 | 8 | 0 | 11 |
| P40261 | NNMT | 29.6 | 0 | 0 | 0 | 0 | 0 | 3 |
| P40429 | RPL13A | 23.6 | 3 | 3 | 3 | 7 | 0 | 10 |
| P40925 | MDH1 | 36.4 | 3 | 3 | 4 | 0 | 0 | 7 |
| P40926 | MDH2 | 35.5 | 0 | 0 | 0 | 0 | 0 | 5 |
| P41091 | EIF2S3 | 51.1 | 0 | 0 | 0 | 0 | 0 | 6 |
| P41250 | GARS1 | 83.1 | 0 | 0 | 0 | 0 | 0 | 6 |
| P41252 | IARS1 | 144.4 | 0 | 0 | 0 | 4 | 0 | 18 |
| P42677 | RPS27 | 9.5 | 0 | 0 | 0 | 0 | 0 | 3 |
| P42766 | RPL35 | 14.5 | 0 | 0 | 0 | 3 | 0 | 4 |
| P42892 | ECE1 | 87.1 | 0 | 0 | 4 | 0 | 0 | 8 |
| P43034 | PAFAH1B1 | 46.6 | 0 | 0 | 0 | 0 | 0 | 7 |
| P43121 | MCAM | 71.6 | 5 | 6 | 8 | 7 | 8 | 18 |
| P43243 | MATR3 | 94.6 | 0 | 0 | 0 | 0 | 0 | 5 |
| P43490 | NAMPT | 55.5 | 0 | 0 | 0 | 0 | 0 | 5 |
| P43686 | PSMC4 | 47.3 | 0 | 0 | 0 | 0 | 0 | 4 |
| P46063 | RECQL | 73.4 | 0 | 0 | 0 | 0 | 0 | 6 |
| P46087 | NOP2 | 89.2 | 0 | 0 | 0 | 0 | 0 | 5 |
| P46776 | RPL27A | 16.6 | 0 | 0 | 0 | 3 | 0 | 6 |
| P46777 | RPL5 | 34.3 | 4 | 4 | 7 | 6 | 0 | 9 |
| P46778 | RPL21 | 18.6 | 0 | 0 | 0 | 3 | 0 | 3 |
| P46779 | RPL28 | 15.7 | 0 | 0 | 0 | 0 | 0 | 3 |
| P46781 | RPS9 | 22.6 | 0 | 3 | 3 | 7 | 3 | 13 |
| P46782 | RPS5 | 22.9 | 0 | 0 | 4 | 0 | 0 | 3 |
| P46940 | IQGAP1 | 189.1 | 12 | 11 | 17 | 25 | 10 | 38 |
| P47755 | CAPZA2 | 32.9 | 0 | 0 | 0 | 0 | 0 | 3 |
| P47756 | CAPZB | 30.6 | 0 | 0 | 0 | 3 | 0 | 4 |
| P47897 | QARS1 | 87.7 | 0 | 0 | 0 | 0 | 0 | 6 |
| P48059 | LIMS1 | 37.2 | 0 | 0 | 0 | 0 | 0 | 3 |
| P48509 | CD151 | 28.3 | 0 | 0 | 0 | 0 | 0 | 3 |
| P48556 | PSMD8 | 39.6 | 0 | 0 | 0 | 0 | 0 | 3 |
| P48643 | CCT5 | 59.6 | 0 | 3 | 4 | 7 | 0 | 11 |
| P49184 | DNASE1L1 | 33.9 | 0 | 0 | 0 | 0 | 0 | 3 |
| P49189 | ALDH9A1 | 53.8 | 3 | 0 | 0 | 0 | 0 | 4 |
| P49327 | FASN | 273.3 | 7 | 0 | 13 | 20 | 3 | 43 |
| P49368 | CCT3 | 60.5 | 4 | 3 | 7 | 5 | 0 | 16 |
| P49588 | AARS1 | 106.7 | 0 | 0 | 0 | 0 | 0 | 5 |
| P49591 | SARS1 | 58.7 | 0 | 0 | 0 | 0 | 0 | 6 |
| P50281 | MMP14 | 65.9 | 0 | 0 | 0 | 0 | 0 | 5 |
| P50395 | GDI2 | 50.6 | 5 | 6 | 9 | 6 | 0 | 8 |
| P50454 | SERPINH1 | 46.4 | 0 | 0 | 0 | 0 | 0 | 3 |
| P50570 | DNM2 | 98 | 0 | 0 | 0 | 0 | 0 | 3 |
| P50914 | RPL14 | 23.4 | 3 | 0 | 4 | 5 | 3 | 5 |
| P50990 | CCT8 | 59.6 | 0 | 3 | 6 | 9 | 0 | 8 |
| P50991 | CCT4 | 57.9 | 3 | 4 | 4 | 7 | 0 | 14 |
| P50995 | ANXA11 | 54.4 | 5 | 7 | 7 | 18 | 11 | 24 |
| P51148 | RAB5C | 23.5 | 3 | 3 | 3 | 0 | 0 | 3 |
| P51149 | RAB7A | 23.5 | 5 | 3 | 4 | 5 | 3 | 11 |
| P51884 | LUM | 38.4 | 0 | 3 | 3 | 0 | 0 | 0 |
| P52209 | PGD | 53.1 | 3 | 0 | 3 | 0 | 0 | 7 |
| P52272 | HNRNPM | 77.5 | 0 | 0 | 0 | 0 | 0 | 6 |
| P52292 | KPNA2 | 57.8 | 0 | 0 | 0 | 0 | 0 | 4 |
| P52565 | ARHGDIA | 23.2 | 3 | 0 | 5 | 4 | 0 | 5 |
| P52566 | ARHGDIB | 23 | 3 | 0 | 3 | 0 | 0 | 6 |
| P52907 | CAPZA1 | 32.9 | 0 | 0 | 3 | 3 | 0 | 0 |
| P53396 | ACLY | 120.8 | 6 | 4 | 9 | 7 | 3 | 19 |
| P53618 | COPB1 | 107.1 | 0 | 0 | 0 | 0 | 0 | 3 |
| P53621 | COPA | 138.3 | 0 | 0 | 3 | 7 | 0 | 16 |
| P53985 | SLC16A1 | 53.9 | 0 | 0 | 0 | 0 | 0 | 3 |
| P53990 | IST1 | 39.7 | 0 | 0 | 4 | 5 | 5 | 12 |
| P54136 | RARS1 | 75.3 | 0 | 0 | 0 | 0 | 0 | 6 |
| P54709 | ATP1B3 | 31.5 | 0 | 0 | 4 | 0 | 0 | 3 |
| P54920 | NAPA | 33.2 | 0 | 0 | 0 | 0 | 0 | 4 |
| P55010 | EIF5 | 49.2 | 0 | 0 | 0 | 0 | 0 | 3 |
| P55011 | SLC12A2 | 131.4 | 0 | 0 | 0 | 0 | 0 | 5 |
| P55060 | CSE1L | 110.3 | 7 | 0 | 3 | 0 | 0 | 4 |
| P55072 | VCP | 89.3 | 19 | 20 | 22 | 8 | 7 | 29 |
| P55209 | NAP1L1 | 45.3 | 0 | 0 | 3 | 0 | 0 | 4 |
| P55786 | NPEPPS | 103.2 | 0 | 0 | 0 | 0 | 0 | 8 |
| P55884 | EIF3B | 92.4 | 0 | 0 | 0 | 0 | 0 | 4 |
| P56192 | MARS1 | 101.1 | 0 | 0 | 4 | 0 | 0 | 0 |
| P56537 | EIF6 | 26.6 | 0 | 0 | 0 | 0 | 0 | 3 |
| P59998 | ARPC4 | 19.7 | 0 | 0 | 0 | 4 | 0 | 3 |
| P60033 | CD81 | 25.8 | 3 | 4 | 4 | 4 | 0 | 7 |
| P60174 | TPI1 | 26.7 | 8 | 9 | 10 | 8 | 6 | 13 |
| P60228 | EIF3E | 52.2 | 0 | 0 | 0 | 0 | 0 | 6 |
| P60660 | MYL6 | 16.9 | 0 | 0 | 0 | 5 | 4 | 4 |
| P60842 | EIF4A1 | 46.1 | 4 | 4 | 0 | 5 | 0 | 5 |
| P60866 | RPS20 | 13.4 | 0 | 3 | 0 | 3 | 0 | 4 |
| P60903 | S100A10 | 11.2 | 0 | 0 | 0 | 0 | 0 | 4 |
| P60953 | CDC42 | 21.2 | 0 | 3 | 3 | 0 | 0 | 3 |
| P61006 | RAB8A | 23.7 | 0 | 0 | 3 | 0 | 0 | 0 |
| P61019 | RAB2A | 23.5 | 0 | 0 | 0 | 0 | 0 | 5 |
| P61026 | RAB10 | 22.5 | 0 | 0 | 0 | 3 | 0 | 3 |
| P61106 | RAB14 | 23.9 | 0 | 0 | 0 | 0 | 0 | 5 |
| P61158 | ACTR3 | 47.3 | 0 | 0 | 3 | 0 | 0 | 8 |
| P61160 | ACTR2 | 44.7 | 0 | 0 | 0 | 3 | 0 | 3 |
| P61163 | ACTR1A | 42.6 | 0 | 0 | 0 | 0 | 0 | 3 |
| P61204 | ARF3 | 20.6 | 5 | 0 | 0 | 4 | 3 | 5 |
| P61221 | ABCE1 | 67.3 | 0 | 0 | 0 | 0 | 0 | 10 |
| P61224 | RAP1B | 20.8 | 5 | 3 | 6 | 4 | 3 | 9 |
| P61225 | RAP2B | 20.5 | 0 | 0 | 0 | 3 | 0 | 7 |
| P61247 | RPS3A | 29.9 | 5 | 5 | 5 | 8 | 4 | 10 |
| P61254 | RPL26 | 17.2 | 3 | 3 | 0 | 0 | 0 | 8 |
| P61313 | RPL15 | 24.1 | 0 | 0 | 0 | 5 | 0 | 6 |
| P61353 | RPL27 | 15.8 | 3 | 0 | 0 | 4 | 0 | 6 |
| P61586 | RHOA | 21.8 | 0 | 0 | 0 | 0 | 3 | 0 |
| P61769 | B2M | 13.7 | 0 | 0 | 0 | 3 | 0 | 0 |
| P61978 | HNRNPK | 50.9 | 0 | 0 | 4 | 6 | 0 | 7 |
| P61981 | YWHAG | 28.3 | 0 | 0 | 0 | 0 | 0 | 3 |
| P62070 | RRAS2 | 23.4 | 0 | 0 | 3 | 0 | 0 | 4 |
| P62081 | RPS7 | 22.1 | 0 | 0 | 3 | 5 | 0 | 6 |
| P62191 | PSMC1 | 49.2 | 0 | 0 | 0 | 0 | 0 | 3 |
| P62195 | PSMC5 | 45.6 | 0 | 0 | 0 | 0 | 0 | 3 |
| P62241 | RPS8 | 24.2 | 7 | 5 | 5 | 7 | 5 | 13 |
| P62244 | RPS15A | 14.8 | 3 | 4 | 3 | 4 | 0 | 8 |
| P62249 | RPS16 | 16.4 | 3 | 4 | 4 | 6 | 5 | 11 |
| P62258 | YWHAE | 29.2 | 0 | 0 | 0 | 6 | 6 | 9 |
| P62263 | RPS14 | 16.3 | 3 | 5 | 5 | 5 | 0 | 7 |
| P62269 | RPS18 | 17.7 | 4 | 3 | 4 | 6 | 5 | 9 |
| P62277 | RPS13 | 17.2 | 6 | 4 | 5 | 6 | 3 | 7 |
| P62280 | RPS11 | 18.4 | 3 | 6 | 5 | 7 | 5 | 11 |
| P62333 | PSMC6 | 44.1 | 0 | 0 | 0 | 0 | 0 | 4 |
| P62424 | RPL7A | 30 | 5 | 5 | 7 | 10 | 5 | 12 |
| P62491 | RAB11A | 24.4 | 3 | 0 | 0 | 4 | 0 | 7 |
| P62701 | RPS4X | 29.6 | 3 | 4 | 6 | 7 | 0 | 11 |
| P62750 | RPL23A | 17.7 | 3 | 0 | 3 | 3 | 0 | 4 |
| P62753 | RPS6 | 28.7 | 4 | 5 | 5 | 5 | 3 | 11 |
| P62805 | H4C16 | 11.4 | 7 | 7 | 8 | 10 | 8 | 13 |
| P62826 | RAN | 24.4 | 5 | 3 | 3 | 3 | 4 | 11 |
| P62829 | RPL23 | 14.9 | 3 | 3 | 3 | 0 | 0 | 4 |
| P62847 | RPS24 | 15.4 | 0 | 0 | 3 | 4 | 0 | 4 |
| P62851 | RPS25 | 13.7 | 3 | 0 | 3 | 3 | 0 | 4 |
| P62873 | GNB1 | 37.4 | 4 | 5 | 5 | 0 | 0 | 10 |
| P62879 | GNB2 | 37.3 | 4 | 3 | 6 | 3 | 0 | 5 |
| P62888 | RPL30 | 12.8 | 4 | 3 | 4 | 4 | 0 | 4 |
| P62899 | RPL31 | 14.5 | 0 | 0 | 0 | 0 | 0 | 7 |
| P62906 | RPL10A | 24.8 | 0 | 3 | 4 | 6 | 0 | 6 |
| P62910 | RPL32 | 15.9 | 0 | 0 | 0 | 0 | 0 | 5 |
| P62913 | RPL11 | 20.2 | 4 | 0 | 0 | 0 | 0 | 4 |
| P62917 | RPL8 | 28 | 0 | 0 | 3 | 3 | 0 | 6 |
| P62937 | PPIA | 18 | 5 | 6 | 7 | 7 | 5 | 9 |
| P62979 | RPS27A | 18 | 6 | 0 | 6 | 5 | 0 | 0 |
| P62987 | UBA52 | 14.7 | 0 | 5 | 0 | 0 | 4 | 10 |
| P63000 | RAC1 | 21.4 | 0 | 0 | 3 | 4 | 0 | 5 |
| P63010 | AP2B1 | 104.5 | 0 | 0 | 0 | 4 | 0 | 0 |
| P63104 | YWHAZ | 27.7 | 3 | 3 | 5 | 7 | 4 | 10 |
| P63173 | RPL38 | 8.2 | 0 | 0 | 0 | 0 | 0 | 3 |
| P63244 | RACK1 | 35.1 | 5 | 5 | 6 | 4 | 0 | 12 |
| P63313 | TMSB10 | 5 | 0 | 0 | 0 | 3 | 0 | 0 |
| P67936 | TPM4 | 28.5 | 0 | 0 | 0 | 4 | 3 | 4 |
| P68104 | EEF1A1 | 50.1 | 10 | 7 | 8 | 10 | 7 | 13 |
| P68366 | TUBA4A | 49.9 | 5 | 3 | 4 | 3 | 0 | 5 |
| P68371 | TUBB4B | 49.8 | 3 | 4 | 3 | 3 | 3 | 3 |
| P68400 | CSNK2A1 | 45.1 | 0 | 0 | 0 | 0 | 0 | 3 |
| P69905 | HBA2 | 15.2 | 4 | 5 | 3 | 4 | 3 | 5 |
| P78371 | CCT2 | 57.5 | 0 | 3 | 5 | 8 | 3 | 13 |
| P78527 | PRKDC | 468.8 | 0 | 0 | 4 | 0 | 0 | 28 |
| P80723 | BASP1 | 22.7 | 0 | 0 | 0 | 5 | 4 | 3 |
| P83731 | RPL24 | 17.8 | 0 | 3 | 5 | 5 | 0 | 7 |
| P84095 | RHOG | 21.3 | 0 | 0 | 0 | 0 | 0 | 6 |
| P84098 | RPL19 | 23.5 | 0 | 0 | 0 | 0 | 0 | 3 |
| P98160 | HSPG2 | 468.5 | 38 | 33 | 48 | 0 | 5 | 49 |
| Q00610 | CLTC | 191.5 | 52 | 58 | 57 | 49 | 31 | 69 |
| Q00839 | HNRNPU | 90.5 | 3 | 4 | 6 | 0 | 0 | 15 |
| Q01082 | SPTBN1 | 274.4 | 0 | 0 | 5 | 0 | 0 | 11 |
| Q01518 | CAP1 | 51.9 | 0 | 3 | 0 | 3 | 0 | 9 |
| Q01813 | PFKP | 85.5 | 5 | 3 | 10 | 11 | 3 | 17 |
| Q01995 | TAGLN | 22.6 | 0 | 0 | 0 | 0 | 0 | 4 |
| Q02241 | KIF23 | 110 | 0 | 0 | 0 | 0 | 0 | 6 |
| Q02413 | DSG1 | 113.7 | 11 | 0 | 8 | 8 | 0 | 8 |
| Q02543 | RPL18A | 20.7 | 0 | 0 | 0 | 3 | 0 | 8 |
| Q02750 | MAP2K1 | 43.4 | 0 | 0 | 0 | 0 | 0 | 3 |
| Q02809 | PLOD1 | 83.5 | 6 | 5 | 9 | 0 | 0 | 9 |
| Q02878 | RPL6 | 32.7 | 5 | 5 | 6 | 12 | 5 | 13 |
| Q02952 | AKAP12 | 191.4 | 0 | 0 | 0 | 0 | 0 | 3 |
| Q03135 | CAV1 | 20.5 | 0 | 0 | 0 | 3 | 0 | 4 |
| Q03405 | PLAUR | 37 | 0 | 0 | 0 | 3 | 0 | 6 |
| Q04446 | GBE1 | 80.4 | 0 | 0 | 0 | 0 | 0 | 3 |
| Q04637 | EIF4G1 | 175.4 | 0 | 0 | 0 | 0 | 0 | 3 |
| Q04695 | KRT17 | 48.1 | 0 | 0 | 0 | 0 | 0 | 3 |
| Q04917 | YWHAH | 28.2 | 0 | 0 | 0 | 0 | 0 | 5 |
| Q06033 | ITIH3 | 99.8 | 3 | 3 | 3 | 0 | 0 | 0 |
| Q06210 | GFPT1 | 78.8 | 0 | 0 | 0 | 0 | 0 | 5 |
| Q06323 | PSME1 | 28.7 | 0 | 0 | 0 | 4 | 0 | 7 |
| Q06830 | PRDX1 | 22.1 | 6 | 5 | 11 | 6 | 3 | 11 |
| Q07020 | RPL18 | 21.6 | 0 | 4 | 4 | 7 | 4 | 7 |
| Q07955 | SRSF1 | 27.7 | 0 | 0 | 0 | 3 | 0 | 6 |
| Q08188 | TGM3 | 76.6 | 3 | 0 | 0 | 4 | 0 | 0 |
| Q08211 | DHX9 | 140.9 | 0 | 0 | 0 | 3 | 0 | 8 |
| Q08380 | LGALS3BP | 65.3 | 6 | 5 | 7 | 9 | 6 | 11 |
| Q08431 | MFGE8 | 43.1 | 7 | 5 | 8 | 4 | 0 | 9 |
| Q08554 | DSC1 | 99.9 | 4 | 0 | 3 | 3 | 0 | 3 |
| Q09666 | AHNAK | 628.7 | 4 | 0 | 7 | 8 | 9 | 24 |
| Q12805 | EFEMP1 | 54.6 | 10 | 11 | 11 | 0 | 0 | 0 |
| Q12846 | STX4 | 34.2 | 0 | 0 | 0 | 0 | 0 | 3 |
| Q12904 | AIMP1 | 34.3 | 0 | 0 | 0 | 0 | 0 | 3 |
| Q12905 | ILF2 | 43 | 0 | 0 | 0 | 0 | 0 | 3 |
| Q13045 | FLII | 144.7 | 0 | 0 | 0 | 0 | 0 | 3 |
| Q13200 | PSMD2 | 100.1 | 0 | 0 | 0 | 3 | 0 | 7 |
| Q13201 | MMRN1 | 138 | 0 | 0 | 0 | 0 | 0 | 4 |
| Q13418 | ILK | 51.4 | 3 | 0 | 3 | 0 | 0 | 3 |
| Q13443 | ADAM9 | 90.5 | 0 | 0 | 0 | 0 | 0 | 12 |
| Q13509 | TUBB3 | 50.4 | 0 | 0 | 3 | 0 | 0 | 4 |
| Q13530 | SERINC3 | 52.5 | 0 | 0 | 0 | 0 | 0 | 4 |
| Q13620 | CUL4B | 103.9 | 0 | 0 | 0 | 0 | 0 | 4 |
| Q13740 | ALCAM | 65.1 | 0 | 0 | 0 | 0 | 0 | 4 |
| Q13813 | SPTAN1 | 284.4 | 0 | 0 | 0 | 5 | 0 | 14 |
| Q13835 | PKP1 | 82.8 | 3 | 0 | 0 | 3 | 0 | 0 |
| Q14152 | EIF3A | 166.5 | 0 | 0 | 3 | 4 | 0 | 7 |
| Q14204 | DYNC1H1 | 532.1 | 11 | 7 | 16 | 17 | 0 | 68 |
| Q14254 | FLOT2 | 47 | 0 | 0 | 0 | 0 | 0 | 8 |
| Q14315 | FLNC | 290.8 | 0 | 0 | 5 | 0 | 0 | 14 |
| Q14697 | GANAB | 106.8 | 0 | 3 | 5 | 0 | 0 | 4 |
| Q14699 | RFTN1 | 63.1 | 3 | 0 | 3 | 0 | 0 | 4 |
| Q14764 | MVP | 99.3 | 16 | 26 | 27 | 31 | 26 | 45 |
| Q14766 | LTBP1 | 186.7 | 0 | 0 | 3 | 0 | 0 | 6 |
| Q14974 | KPNB1 | 97.1 | 4 | 0 | 5 | 0 | 0 | 5 |
| Q15008 | PSMD6 | 45.5 | 0 | 0 | 0 | 3 | 0 | 6 |
| Q15019 | SEPTIN2 | 41.5 | 0 | 0 | 3 | 4 | 0 | 7 |
| Q15063 | POSTN | 93.3 | 0 | 4 | 10 | 0 | 0 | 0 |
| Q15149 | PLEC | 531.5 | 0 | 0 | 0 | 3 | 0 | 13 |
| Q15286 | RAB35 | 23 | 0 | 0 | 0 | 0 | 0 | 3 |
| Q15365 | PCBP1 | 37.5 | 0 | 0 | 3 | 0 | 0 | 5 |
| Q15366 | PCBP2 | 38.6 | 0 | 0 | 3 | 0 | 0 | 3 |
| Q15435 | PPP1R7 | 41.5 | 0 | 0 | 0 | 0 | 0 | 3 |
| Q15582 | TGFBI | 74.6 | 4 | 6 | 9 | 4 | 0 | 10 |
| Q15758 | SLC1A5 | 56.6 | 3 | 4 | 3 | 0 | 0 | 4 |
| Q15833 | STXBP2 | 66.4 | 0 | 0 | 0 | 0 | 0 | 3 |
| Q16181 | SEPTIN7 | 50.6 | 0 | 0 | 4 | 0 | 0 | 5 |
| Q16222 | UAP1 | 58.7 | 0 | 0 | 0 | 0 | 0 | 3 |
| Q16363 | LAMA4 | 202.4 | 5 | 0 | 7 | 0 | 0 | 11 |
| Q16401 | PSMD5 | 56.2 | 0 | 0 | 0 | 0 | 0 | 4 |
| Q16555 | DPYSL2 | 62.3 | 0 | 0 | 0 | 0 | 0 | 10 |
| Q16563 | SYPL1 | 28.5 | 0 | 0 | 0 | 0 | 0 | 3 |
| Q16658 | FSCN1 | 54.5 | 4 | 3 | 7 | 3 | 3 | 8 |
| Q16777 | H2AC20 | 14 | 0 | 0 | 0 | 3 | 0 | 0 |
| Q16819 | MEP1A | 84.4 | 0 | 0 | 0 | 0 | 0 | 3 |
| Q16851 | UGP2 | 56.9 | 0 | 0 | 0 | 0 | 0 | 6 |
| Q16881 | TXNRD1 | 70.9 | 0 | 0 | 0 | 0 | 0 | 7 |
| Q32P51 | HNRNPA1L2 | 34.2 | 0 | 0 | 0 | 3 | 0 | 0 |
| Q53EZ4 | CEP55 | 54.1 | 0 | 0 | 0 | 0 | 0 | 10 |
| Q5D862 | FLG2 | 247.9 | 0 | 0 | 0 | 3 | 0 | 0 |
| Q5JWF2 | GNAS | 111 | 0 | 0 | 0 | 3 | 0 | 8 |
| Q5QNW6 | H2BC18 | 13.9 | 3 | 3 | 0 | 0 | 3 | 0 |
| Q5SSJ5 | HP1BP3 | 61.2 | 0 | 0 | 0 | 3 | 0 | 7 |
| Q5T749 | KPRP | 64.1 | 5 | 0 | 0 | 4 | 0 | 5 |
| Q5VW32 | BROX | 46.4 | 0 | 0 | 0 | 0 | 0 | 4 |
| Q6DD88 | ATL3 | 60.5 | 0 | 0 | 0 | 0 | 0 | 3 |
| Q6IS14 | EIF5AL1 | 16.8 | 0 | 0 | 0 | 0 | 0 | 3 |
| Q6KB66 | KRT80 | 50.5 | 3 | 0 | 0 | 3 | 0 | 0 |
| Q6NZI2 | CAVIN1 | 43.5 | 0 | 0 | 3 | 3 | 3 | 6 |
| Q6PIU2 | NCEH1 | 45.8 | 0 | 0 | 0 | 0 | 0 | 3 |
| Q6UVK1 | CSPG4 | 250.4 | 0 | 0 | 0 | 0 | 0 | 21 |
| Q6YHK3 | CD109 | 161.6 | 0 | 0 | 0 | 0 | 0 | 3 |
| Q71U36 | TUBA1A | 50.1 | 0 | 0 | 0 | 5 | 0 | 0 |
| Q7KZF4 | SND1 | 101.9 | 0 | 0 | 0 | 0 | 0 | 7 |
| Q7L1Q6 | BZW1 | 48 | 0 | 0 | 0 | 0 | 0 | 4 |
| Q7L2H7 | EIF3M | 42.5 | 0 | 0 | 0 | 0 | 0 | 3 |
| Q7L576 | CYFIP1 | 145.1 | 0 | 0 | 0 | 0 | 0 | 10 |
| Q7Z2W4 | ZC3HAV1 | 101.4 | 0 | 0 | 0 | 0 | 0 | 3 |
| Q7Z7G0 | ABI3BP | 117.8 | 0 | 0 | 0 | 0 | 0 | 3 |
| Q86UX7 | FERMT3 | 75.9 | 0 | 0 | 0 | 0 | 0 | 5 |
| Q86VP6 | CAND1 | 136.3 | 3 | 0 | 0 | 0 | 0 | 7 |
| Q86Y82 | STX12 | 31.6 | 0 | 0 | 0 | 0 | 0 | 4 |
| Q86YZ3 | HRNR | 282.2 | 5 | 0 | 0 | 3 | 0 | 3 |
| Q8IVF7 | FMNL3 | 117.1 | 0 | 0 | 0 | 0 | 0 | 3 |
| Q8IWA5 | SLC44A2 | 80.1 | 0 | 0 | 0 | 0 | 0 | 4 |
| Q8IWT6 | LRRC8A | 94.1 | 0 | 0 | 0 | 0 | 0 | 5 |
| Q8IX04 | UEVLD | 52.2 | 0 | 0 | 0 | 0 | 0 | 4 |
| Q8IZ83 | ALDH16A1 | 85.1 | 7 | 7 | 4 | 0 | 0 | 5 |
| Q8IZL8 | PELP1 | 119.6 | 0 | 0 | 0 | 0 | 0 | 7 |
| Q8IZP2 | ST13P4 | 27.4 | 0 | 0 | 3 | 3 | 0 | 3 |
| Q8N1N4 | KRT78 | 56.8 | 6 | 0 | 4 | 0 | 3 | 3 |
| Q8N5I2 | ARRDC1 | 46 | 0 | 0 | 0 | 0 | 0 | 5 |
| Q8N699 | MYCT1 | 26.6 | 0 | 0 | 3 | 0 | 0 | 5 |
| Q8N9N7 | LRRC57 | 26.7 | 0 | 0 | 0 | 0 | 0 | 5 |
| Q8NG11 | TSPAN14 | 30.7 | 0 | 0 | 0 | 0 | 0 | 8 |
| Q8WUM4 | PDCD6IP | 96 | 13 | 14 | 21 | 20 | 18 | 50 |
| Q8WV92 | MITD1 | 29.3 | 0 | 0 | 0 | 0 | 0 | 4 |
| Q8WWI5 | SLC44A1 | 73.3 | 0 | 0 | 0 | 0 | 0 | 9 |
| Q92499 | DDX1 | 82.4 | 0 | 0 | 0 | 0 | 0 | 5 |
| Q92542 | NCSTN | 78.4 | 0 | 0 | 0 | 0 | 0 | 4 |
| Q92626 | PXDN | 165.2 | 17 | 16 | 29 | 9 | 0 | 18 |
| Q92692 | NECTIN2 | 57.7 | 0 | 0 | 0 | 0 | 0 | 4 |
| Q92743 | HTRA1 | 51.3 | 0 | 0 | 3 | 0 | 0 | 5 |
| Q92841 | DDX17 | 80.2 | 0 | 0 | 0 | 4 | 0 | 11 |
| Q92882 | OSTF1 | 23.8 | 0 | 0 | 0 | 0 | 0 | 3 |
| Q92930 | RAB8B | 23.6 | 0 | 0 | 0 | 0 | 0 | 4 |
| Q92945 | KHSRP | 73.1 | 0 | 0 | 0 | 0 | 0 | 3 |
| Q93050 | ATP6V0A1 | 96.4 | 0 | 0 | 0 | 0 | 0 | 3 |
| Q969P0 | IGSF8 | 65 | 0 | 0 | 0 | 0 | 0 | 7 |
| Q969Q0 | RPL36AL | 12.5 | 0 | 0 | 0 | 0 | 0 | 3 |
| Q96AC1 | FERMT2 | 77.8 | 0 | 0 | 0 | 0 | 0 | 9 |
| Q96CW1 | AP2M1 | 49.6 | 0 | 0 | 0 | 0 | 0 | 7 |
| Q96CX2 | KCTD12 | 35.7 | 0 | 0 | 0 | 0 | 0 | 7 |
| Q96F07 | CYFIP2 | 148.3 | 0 | 0 | 4 | 0 | 0 | 0 |
| Q96FN4 | CPNE2 | 61.2 | 3 | 3 | 3 | 0 | 0 | 9 |
| Q96P63 | SERPINB12 | 46.2 | 5 | 0 | 0 | 5 | 0 | 3 |
| Q96QK1 | VPS35 | 91.6 | 0 | 0 | 0 | 0 | 0 | 6 |
| Q96QV1 | HHIP | 78.8 | 9 | 7 | 12 | 0 | 0 | 14 |
| Q96S97 | MYADM | 35.3 | 0 | 0 | 0 | 0 | 0 | 3 |
| Q96TA1 | NIBAN2 | 84.1 | 0 | 0 | 3 | 5 | 0 | 11 |
| Q99460 | PSMD1 | 105.8 | 0 | 0 | 0 | 0 | 0 | 3 |
| Q99497 | PARK7 | 19.9 | 0 | 0 | 0 | 0 | 0 | 4 |
| Q99536 | VAT1 | 41.9 | 3 | 4 | 4 | 0 | 0 | 5 |
| Q99808 | SLC29A1 | 50.2 | 0 | 0 | 3 | 0 | 0 | 3 |
| Q99816 | TSG101 | 43.9 | 0 | 0 | 0 | 0 | 0 | 8 |
| Q99829 | CPNE1 | 59 | 0 | 0 | 0 | 0 | 0 | 9 |
| Q99832 | CCT7 | 59.3 | 0 | 0 | 5 | 4 | 0 | 13 |
| Q99873 | PRMT1 | 42.4 | 0 | 0 | 0 | 0 | 0 | 4 |
| Q99878 | H2AC14 | 13.9 | 4 | 0 | 0 | 0 | 0 | 0 |
| Q99961 | SH3GL1 | 41.5 | 0 | 0 | 0 | 0 | 0 | 8 |
| Q99985 | SEMA3C | 85.2 | 3 | 0 | 7 | 0 | 0 | 3 |
| Q9BQE3 | TUBA1C | 49.9 | 0 | 0 | 0 | 0 | 5 | 8 |
| Q9BSJ8 | ESYT1 | 122.8 | 0 | 0 | 0 | 0 | 0 | 3 |
| Q9BUF5 | TUBB6 | 49.8 | 4 | 0 | 6 | 0 | 0 | 6 |
| Q9BV38 | WDR18 | 47.4 | 0 | 0 | 0 | 0 | 0 | 5 |
| Q9BWD1 | ACAT2 | 41.3 | 0 | 0 | 0 | 0 | 0 | 5 |
| Q9BXJ9 | NAA15 | 101.2 | 0 | 0 | 0 | 0 | 0 | 3 |
| Q9BY44 | EIF2A | 64.9 | 0 | 0 | 0 | 0 | 0 | 3 |
| Q9BY76 | ANGPTL4 | 45.2 | 0 | 0 | 3 | 0 | 0 | 3 |
| Q9C0H2 | TTYH3 | 57.5 | 0 | 0 | 0 | 0 | 0 | 5 |
| Q9GZM7 | TINAGL1 | 52.4 | 0 | 0 | 6 | 0 | 0 | 3 |
| Q9H0H5 | RACGAP1 | 71 | 0 | 0 | 0 | 0 | 0 | 5 |
| Q9H0U4 | RAB1B | 22.2 | 0 | 0 | 3 | 0 | 0 | 0 |
| Q9H223 | EHD4 | 61.1 | 6 | 5 | 7 | 6 | 0 | 18 |
| Q9H444 | CHMP4B | 24.9 | 0 | 0 | 0 | 0 | 0 | 5 |
| Q9H4B7 | TUBB1 | 50.3 | 4 | 0 | 3 | 0 | 0 | 3 |
| Q9H4G4 | GLIPR2 | 17.2 | 0 | 0 | 3 | 0 | 0 | 6 |
| Q9H4M9 | EHD1 | 60.6 | 5 | 0 | 5 | 4 | 0 | 18 |
| Q9H5V8 | CDCP1 | 92.9 | 0 | 0 | 0 | 0 | 0 | 3 |
| Q9HD42 | CHMP1A | 21.7 | 0 | 0 | 0 | 0 | 0 | 3 |
| Q9NP72 | RAB18 | 23 | 0 | 0 | 0 | 0 | 0 | 3 |
| Q9NQC3 | RTN4 | 129.9 | 0 | 0 | 0 | 4 | 3 | 5 |
| Q9NR30 | DDX21 | 87.3 | 0 | 0 | 0 | 0 | 0 | 5 |
| Q9NR31 | SAR1A | 22.4 | 0 | 0 | 3 | 0 | 0 | 0 |
| Q9NR45 | NANS | 40.3 | 0 | 0 | 0 | 4 | 0 | 6 |
| Q9NRQ2 | PLSCR4 | 37 | 0 | 0 | 0 | 0 | 0 | 3 |
| Q9NRY6 | PLSCR3 | 31.6 | 0 | 0 | 0 | 4 | 3 | 8 |
| Q9NTK5 | OLA1 | 44.7 | 0 | 0 | 0 | 0 | 0 | 3 |
| Q9NXF1 | TEX10 | 105.6 | 0 | 0 | 0 | 0 | 0 | 5 |
| Q9NZM1 | MYOF | 234.6 | 9 | 8 | 20 | 13 | 10 | 55 |
| Q9NZN4 | EHD2 | 61.1 | 0 | 0 | 7 | 3 | 3 | 16 |
| Q9NZT1 | CALML5 | 15.9 | 0 | 0 | 0 | 3 | 0 | 0 |
| Q9NZZ3 | CHMP5 | 24.6 | 0 | 0 | 0 | 0 | 0 | 5 |
| Q9P258 | RCC2 | 56 | 0 | 0 | 0 | 0 | 0 | 3 |
| Q9P265 | DIP2B | 171.4 | 0 | 0 | 0 | 0 | 0 | 14 |
| Q9P273 | TENM3 | 300.8 | 0 | 0 | 0 | 0 | 0 | 6 |
| Q9P2B2 | PTGFRN | 98.5 | 0 | 0 | 4 | 0 | 0 | 7 |
| Q9P2J5 | LARS1 | 134.4 | 0 | 0 | 0 | 0 | 0 | 5 |
| Q9UBI6 | GNG12 | 8 | 0 | 0 | 0 | 0 | 0 | 3 |
| Q9UBV8 | PEF1 | 30.4 | 0 | 0 | 0 | 3 | 0 | 3 |
| Q9UDY2 | TJP2 | 133.9 | 0 | 0 | 0 | 0 | 0 | 3 |
| Q9UDY4 | DNAJB4 | 37.8 | 0 | 0 | 0 | 0 | 0 | 8 |
| Q9UHD8 | SEPTIN9 | 65.4 | 0 | 0 | 0 | 3 | 0 | 4 |
| Q9UHF1 | EGFL7 | 29.6 | 0 | 0 | 0 | 0 | 0 | 5 |
| Q9UHN6 | CEMIP2 | 154.3 | 0 | 0 | 0 | 0 | 0 | 7 |
| Q9UK41 | VPS28 | 25.4 | 0 | 0 | 0 | 0 | 0 | 6 |
| Q9UKK3 | PARP4 | 192.5 | 9 | 6 | 10 | 3 | 0 | 17 |
| Q9UL42 | PNMA2 | 41.5 | 0 | 0 | 0 | 0 | 0 | 6 |
| Q9UL46 | PSME2 | 27.4 | 0 | 0 | 0 | 0 | 0 | 3 |
| Q9ULV4 | CORO1C | 53.2 | 0 | 0 | 3 | 0 | 0 | 10 |
| Q9UMS4 | PRPF19 | 55.1 | 0 | 0 | 0 | 0 | 0 | 4 |
| Q9UN37 | VPS4A | 48.9 | 0 | 0 | 0 | 0 | 0 | 5 |
| Q9UNF0 | PACSIN2 | 55.7 | 0 | 0 | 0 | 0 | 0 | 5 |
| Q9UNH7 | SNX6 | 46.6 | 0 | 0 | 0 | 0 | 0 | 3 |
| Q9UNM6 | PSMD13 | 42.9 | 0 | 0 | 0 | 3 | 0 | 5 |
| Q9UQ80 | PA2G4 | 43.8 | 0 | 0 | 0 | 0 | 0 | 3 |
| Q9Y230 | RUVBL2 | 51.1 | 0 | 0 | 4 | 4 | 0 | 7 |
| Q9Y262 | EIF3L | 66.7 | 0 | 0 | 0 | 4 | 0 | 4 |
| Q9Y265 | RUVBL1 | 50.2 | 0 | 0 | 0 | 4 | 0 | 8 |
| Q9Y2A7 | NCKAP1 | 128.7 | 0 | 0 | 0 | 0 | 0 | 5 |
| Q9Y2J2 | EPB41L3 | 120.6 | 0 | 0 | 3 | 0 | 0 | 3 |
| Q9Y315 | DERA | 35.2 | 5 | 7 | 4 | 3 | 0 | 9 |
| Q9Y3A5 | SBDS | 28.7 | 0 | 0 | 0 | 0 | 0 | 5 |
| Q9Y3I0 | RTCB | 55.2 | 0 | 0 | 0 | 0 | 0 | 4 |
| Q9Y3U8 | RPL36 | 12.2 | 0 | 0 | 0 | 3 | 3 | 4 |
| Q9Y490 | TLN1 | 269.6 | 30 | 15 | 33 | 14 | 6 | 43 |
| Q9Y4K0 | LOXL2 | 86.7 | 0 | 0 | 5 | 0 | 0 | 6 |
| Q9Y4W2 | LAS1L | 83 | 0 | 0 | 0 | 0 | 0 | 4 |
| Q9Y5B9 | SUPT16H | 119.8 | 0 | 0 | 0 | 0 | 0 | 3 |
| Q9Y5K6 | CD2AP | 71.4 | 0 | 0 | 0 | 0 | 0 | 4 |
| Q9Y5P6 | GMPPB | 39.8 | 0 | 0 | 0 | 0 | 0 | 3 |
| Q9Y624 | F11R | 32.6 | 0 | 0 | 0 | 0 | 0 | 5 |
| Q9Y678 | COPG1 | 97.7 | 0 | 0 | 0 | 0 | 0 | 4 |
| Q9Y696 | CLIC4 | 28.8 | 5 | 4 | 11 | 4 | 0 | 9 |
| Q9Y6G9 | DYNC1LI1 | 56.5 | 0 | 0 | 0 | 0 | 0 | 3 |

*Supplementary Table 3. Complete mass spectrometry results: total unique peptides detected per protein across UC Bulk (triplicates) and UC+SEC FR1 (triplicates) preparations.*
